# Monosodium Urate Crystals regulate a unique JNK-dependent macrophage metabolic and inflammatory response

**DOI:** 10.1101/2021.04.14.439881

**Authors:** Isidoro Cobo, Anyan Cheng, Jessica Murillo-Saich, Roxana Coras, Alyssa Torres, Addison J. Lana, Johannes Schlachetzki, Ru Liu-Bryan, Robert Terkeltaub, Elsa Sanchez-Lopez, Christopher K. Glass, Monica Guma

**Affiliations:** Department of Cellular and Molecular Medicine, University of California San Diego, San Diego, La Jolla, CA, USA; Division of Rheumatology, Allergy and Immunology. UCSD School of Medicine, 9500 Gilman Drive, La Jolla, CA, USA; Department of Medicine, Autonomous University of Barcelona, Plaça Cívica, 08193 Bellaterra, Barcelona, Spain; VA San Diego Healthcare System, 3350 La Jolla Village Drive, San Diego, CA, USA; Department of Orthopedic Surgery, University of California San Diego, San Diego, La Jolla, CA, USA

## Abstract

How macrophages are programmed to respond to monosodium urate crystals (MSUc) is incompletely understood partly due to the use of a toll-like receptor-induced priming step. Here, using genome wide transcriptomic analysis and biochemical assays we demonstrate that MSUc alone induces an *in vitro* metabolic-inflammatory transcriptional program in both human and murine macrophages markedly distinct from that induced by LPS. Genes uniquely up-regulated in response to MSUc belonged to lipids, glycolysis, and transport of small molecules via SLC transporters pathways. Sera from individuals and mice with acute gouty arthritis provided further evidence for this metabolic rewiring. This distinct macrophage activation may explain the initiating mechanisms in acute gout flares and is regulated through JUN binding to the promoter of target genes through activation of JNK –but not by P38-in a process that is independent of inflammasome activation. Finally, pharmacological JNK inhibition limited MSUc-induced inflammation in animal models of acute gouty inflammation.

## INTRODUCTION

Macrophages are the primary immune cells displaying an inflammatory response that is intended to restore tissue homeostasis after pathogen infection or tissue damage (1-3). A key feature of macrophages supporting their regulation of inflammatory responses, immunity, tissue homeostasis and repair, is their high degree of plasticity in response to various microenvironmental stimuli (4, 5). However, de-regulated or persistent, inflammation can promote irreversible tissue damage by shifting macrophages phenotypes. Therefore, understanding the mechanisms that govern macrophage biology will be critical to effectively altering their functions for therapeutic purposes by blocking an unwanted pathway or reprogramming macrophages to phenotypes necessary for tissue homeostasis and repair responses (6).

Metabolic reprograming is a hallmark of macrophage activation (7). Recent advances in transcriptomic and metabolomic studies have highlighted the link between metabolic rewiring of macrophages and their functional plasticity (8-10). Many factors in the cell microenvironment activate key signaling pathways that modulate cellular metabolism to support macrophage differentiation and polarization. Indeed, pro-inflammatory macrophages utilize glycolysis and the pentose phosphate pathway (PPP) to meet their ATP requirements, whereas oxidative phosphorylation (OXPHOS) as well as the fatty acid oxidation (FAO) pathways are downregulated. In contrast, in macrophages with anti-inflammatory properties, the Krebs cycle is intact, and their metabolic activity is characterized by enhanced amino acid uptake, enhanced FAO and OXPHOS. These metabolic changes appear to be critical in the pathogenesis of inflammatory and autoimmune diseases (11-13).

Gout is currently the most common cause of inflammatory arthritis. In gout, arthritis and bursitis are caused by tissue deposition of monosodium urate crystals (MSUc) (14, 15). Macrophages, both specialized tissue-resident macrophages and derived from circulating monocytes, are thought to be important for initiating and driving the early proinflammatory phase of acute gout (16-18). The study of the underlying mechanisms of gouty inflammation has yielded some insights into the control of pro-inflammatory cytokines, especially of IL-1β, after macrophage priming with TLR agonists, and inflammasome activation (18, 19), secretion of other cytokines including IL-8, and activation of signaling pathways such as mitogen activated protein kinase (MAPK) and nuclear factor kappa B (20, 21). Of interest, Renaudin et al, recently described a role of glucose metabolism during the inflammatory IL-1β response induced by MSUc in primed-THP1 cells, suggesting that metabolic changes in macrophages after MSUc activation are relevant in gout (22).

Since IL-1β plays an essential role in the pathogenesis of gout flares (18, 19), nearly all mechanistic studies of MSUc-induced macrophage activation *in vitro* introduce a TLR-induced priming step. This results in a robust TLR-dependent transcriptional program and high levels of pro-IL1β that are processed by the inflammasome for secretion. Yet, the inflammatory and metabolic effect that MSUc alone induce in non-primed primary macrophages and the mechanism by which macrophages respond to MSUc and activates this inflammatory and metabolic program is incompletely understood. While TLR-dependent signaling has been linked to the pathophysiology of gout, MSU crystals induce inflammation *in vivo* without prior priming, raising the possibility of an initial cell-autonomous phase that is TLR-independent. To investigate this possibility, we evaluated the transcriptional profile and its regulation in non-primed human monocytes derived macrophages (mDM) and murine bone marrow derived macrophages (BMDM) after MSU crystals alone. Our results indicate that MSUc alone induce a distinct transcriptional program, with a unique inflammatory and metabolic profile, and that up-regulation of JUN and p-JUN^Ser63^ via JNK is necessary for up-regulation of the inflammatory and metabolic transcriptional program induced by MSUc in both mDMs and BMDMs.

## RESULTS

### MSUc induces a distinct transcriptional program in both human and mouse non-primed macrophages

To address the genome-wide contribution of MSUc in non-primed macrophages, we performed RNA sequencing (RNA-Seq) of macrophages differentiated from healthy human monocytes (mDM) and of murine bone marrow derived macrophages (BMDM) stimulated with MSUc (250ug/ml) or LPS (100ng/ml) for 5h. Principal component analysis (PC) highlights the divergence of mDM and BMDM treated with MSUc from untreated cells or cells treated with LPS (**Figure 1A)**. RNA-Seq data was validated by qRT-qPCR in an independent series of experiments (data not shown). Gene set enrichment analysis (GSEA) show those up-regulated genes by MSUc vs. PBS (1480 in mDM or 1366 in BMDM) belong to pathways related to inflammation, metabolism (including lipid metabolism, transport of small molecules, metabolism of carbohydrates and amino acids) and circadian clock (**Figure 1B)** highlighting the importance of MSUc in the regulation of inflammatory and metabolic changes in macrophages.

**Figure 1.**
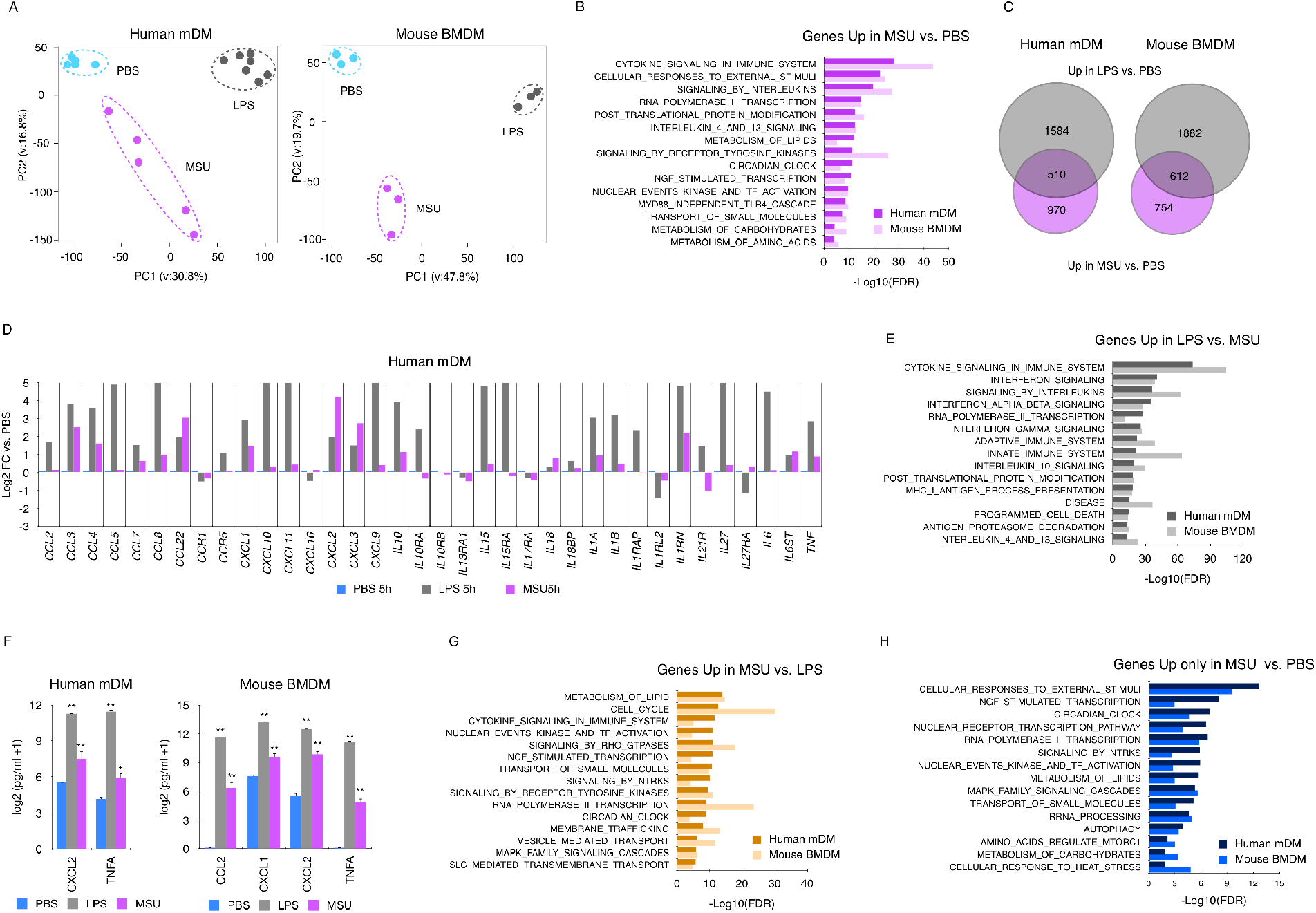
MSUc activates a distinct transcriptional program. (A) Principal component analysis of human mDM (left) or mouse BMDM (right) treated with LPS or MSUc for 5h showing the divergence of the transcriptomic program (n ≥ 5 donors in mDM and n=3 for BMDM). (B) GSEA analysis of genes up-regulated in macrophages treated with MSUc vs. PBS showing inflammatory gene-sets as well as activation of nuclear receptor, NGF, circadian clock and metabolic signaling pathways. (C) Venn diagram showing the overlap between genes up-regulated in macrophages treated with LPS or MSUc. (D) Examples of inflammatory genes up-regulated in mDM treated with LPS or MSUc showing up-regulation of some genes in MSUc but greater up-regulation in LPS. (E) GSEA analysis of genes down-regulated in MSUc vs. LPS showing inflammatory gene-sets including signaling by interferons. (F) ELISA for CXCL2 and TNFA in mDM and additionally CCL2 and CXCL1 for BMDM in the supernatant of macrophages treated with LPS or MSUc showing increased concentration. (G) GSEA using REACTOME of genes up-regulated in MSUc vs. LPS showing metabolic gene sets as well as activation of NGF, receptors tyrosine kinase (RTKs) and circadian clock signaling pathways. (H) GSEA using REACTOME of genes uniquely up-regulated by MSUc showing enrichment in genes sets of response to external stimuli activation of circadian clock, NGF and metabolic signaling pathways. (#=P<0.10;*=P<0.05; **=P<0.01).

We then compared the transcriptional program altered by MSUc with LPS. More than 4000 genes were >2-fold differentially regulated (FDR <0.05) in comparing MSUc-treated BMDMs with LPS-treated BMDMs. Similar differences were observed in mDM, suggesting a distinct macrophage profile after MSUc stimulation (**Supplementary Figure 1A,B)**. Thirty four percent (510/1480) in BMDM and 44% (612/1366) in hMD of genes up-regulated by MSUc vs. PBS were similarly up-regulated by LPS vs. PBS. (**Figure 1C)**. These genes belong mainly to inflammatory signaling pathways **(Supplementary Figure 1C)** including “CCLs”, “CXCLs” and interleukin genes **(Figure 1D and Supplementary Figure 1D)**. As expected, the degree of up-regulation of inflammatory transcripts in LPS is greater than in MSUc (**Figure 1D,E)** and this was accompanied by higher secretion of inflammatory cytokines including CXCL2 and TNFA in mDM and BMDM and CCL2, CXCL1 in BMDM **(Figure 1F)**. Of note, activation of interferon stimulated genes including *CXCL10, IFIT*s and *OAS*s genes were only up-regulated in LPS **(Supplementary Figure 1E,F)**. Interestingly, a set of genes that were up-regulated in MSUc vs. PBS or uniquely up-regulated in MSUc vs. PBS belonged to metabolic pathways, including metabolism of lipids, carbohydrates and amino acids, and transport of small molecules via SLC transporters **(Figure 1G,H)**. These data suggest that MSUc activates a transcriptional program that is in a fashion similar to LPS but distinct in the higher activation of metabolic genes.

### MSUc induces an altered metabolic profile both local and systemically characterized by activation of glycolysis and metabolism of amino acids

The preferential up-regulation of metabolic signaling pathways led us to investigate the effect of MSUc in altering the metabolic profile of macrophages. RNA-Seq showed de-regulation of several transcripts involved in metabolism of carbohydrates in macrophages treated with MSUc or LPS that were validated using RT-qPCR (**Supplementary Figure 2A)**, including up-regulation of *SLC2A1* and *PFKBP3* mRNA **(Figure 2A)**. GLUT1, a glucose transporter encoded by SLC2A1, was observed at protein level using IF **(Figure 2B)** and WB (**Supplementary Figure 2B**) in both BMDMs and mDMs treated with either MSUc or LPS. In addition, we performed one-dimension nuclear magnetic resonance (1D ^1^H-NMR) in the culture media and cell pellet of BMDM and in the supernatant of mDM at 4h and 8h after MSUc or LPS treatment. We found increased concentration of several metabolites related to glucose metabolism including acetate, citrate, formate, fumarate, lactate and succinate in the pellet and supernatant of BMDM treated with MSUc but not LPS (except of increased lactate in BMDM) **(Figure 2C,D, respectively)**. Some of these metabolites also showed higher concentration in the supernatant of mDM treated with MSUc including acetate, citrate and lactate and reduced concentration of succinate **(Figure 2E)**. In addition to altered expression of genes involved in glucose metabolism, we found up-regulation of genes related to amino acid transport in macrophages treated specially with MSUc including *SLC*s transporters such as *SLC3A2, SLC1A4, SLC1A5* and *SLC38A2*, and genes associated with metabolism of amino acids, including genes involved in glycine, serine and threonine metabolism **(Figure 2F)**. These changes were accompanied by increased concentration in the supernatant and pellet of alanine, glutamate, glycine, phenylalanine, threonine and tryptophan of BMDM treated with MSUc but not LPS **(Figure 2G,H, respectively)**. Some of these changes including increased concentration of glutamate and glycine was also observed in mDM treated with MSUc; changes in concentration of other amino acids such as aspartate, glutamine, leucine and isoleucine was also observed in mDM treated with MSUc but did not reach significance in BMDM **(Figure 2I)**. These data indicate that MSUc alters the glucose and amino acid profile of macrophages to a greater extent than LPS.

**Figure 2.**
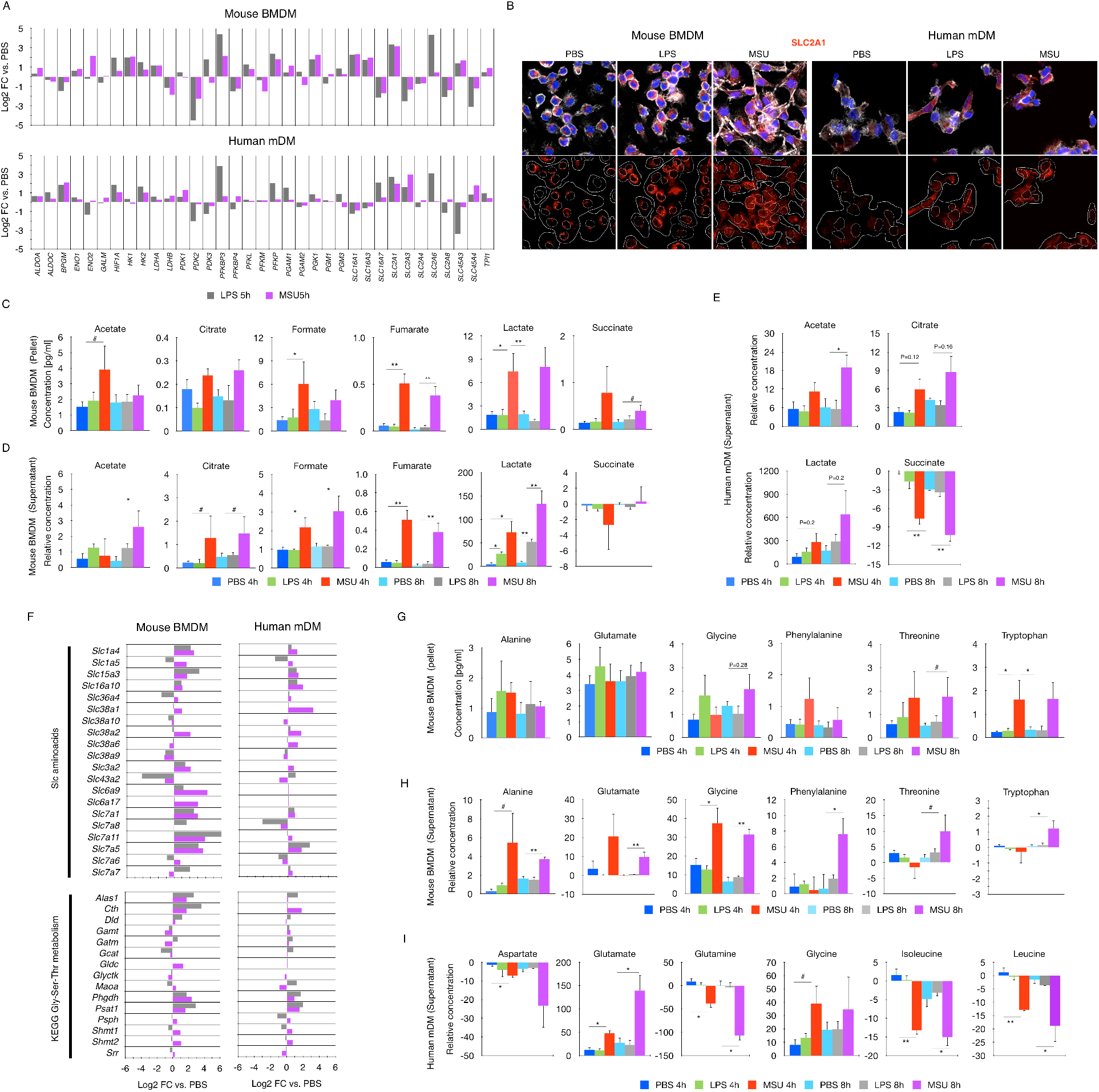
MSUc alters the metabolic program of macrophages. (A) Bar plot showing up-regulation of genes involved in glucose metabolism in macrophages treated with MSUc or LPS. (B) Protein analysis by IF showing up-regulation of SLC2A1 in macrophages treated with MSUc or LPS at 5h. (C-D) Analysis of the concentration of glycolytic metabolites by 1D ^1^H-NMR in the pellet (C) or supernatant (D) or BMDM treated with LPS or MSUc for 4h or 8h showing activation of glycolysis in macrophages treated with MSUc to a higher degree than LPS (n≥4/condition). (E) 1D ^1^H-NMR showing increased levels of acetate, citrate and lactate and reduced levels of succinate in the supernatant of mDM treated with MSUc for 4h or 8h (n=3 donors/condition). (F) Bar graphs showing up-regulation of several amino acid transporters (top) or genes involved in glycine/serine/threonine metabolism (bottom) in macrophages treated with MSUc or LPS for 5h assessed by RNA-Seq. (G,H) 1D ^1^H-NMR showing increased levels of glycine, threonine and tryptophan in the pellet (G) and alanine, glutamate, phenylalanine and tryptophan in supernatant (H) of BMDM treated with MSUc or LPS for 4h or 8h (n≥4/condition). (I) 1D ^1^H-NMR showing increased levels of glutamate and glycine, and reduced levels of aspartate, glutamine, isoleucine and leucine in the supernatant of mDM treated with MSUc for 4h or 8h (n=3 donors/condition). (#=P<0.10;*=P<0.05; **=P<0.01).

We next investigated whether MSUc activates these metabolic pathways *in vivo*. We first explored whether metabolism was altered in a murine air pouch model of acute gouty inflammation. Partial least squares discriminant analysis (PLS-DA) shows the difference metabolic profile of the air pouch lavage of the mice injected with MSUc as assessed by 1D ^1^H-NMR (**Supplementary Figure 3A)**. These changes include reduced concentration of glucose, trimethylamine, alanine, methionine, tyrosine, phenylalanine and an increase of dimethylsulfone **(Supplementary Figure 3B)**. Next, we determined whether local MSUc injection could induced an altered systemic metabolic profile. PLS-DA separates the sera 1D ^1^H-NMR signal into two groups according to mice injected with PBS or with MSUc in the air pouch **(Figure 3A)**. The two most important metabolites to discriminate between groups were acetate and trimethylamine (TMA) and other changes in the metabolic profile include reduced concentration of citrate, formate, fumarate, glucose and lactate with MSUc **(Figure 3B,C)**. Similar to the air pouch gout model in mice, we found distinct metabolic profile of serum of patients suffering an acute gout flare compared with individuals with hyperuricemia (HU) but without current flare (21) including reduced levels of glucose, glutamine, trimethylamine-N-oxide (TMAO), and phosphocholine and increased of dimethylamine **(Figure 3D-F)**. Of interest, glucose was one of metabolites that separated both groups the most.

**Figure 3.**
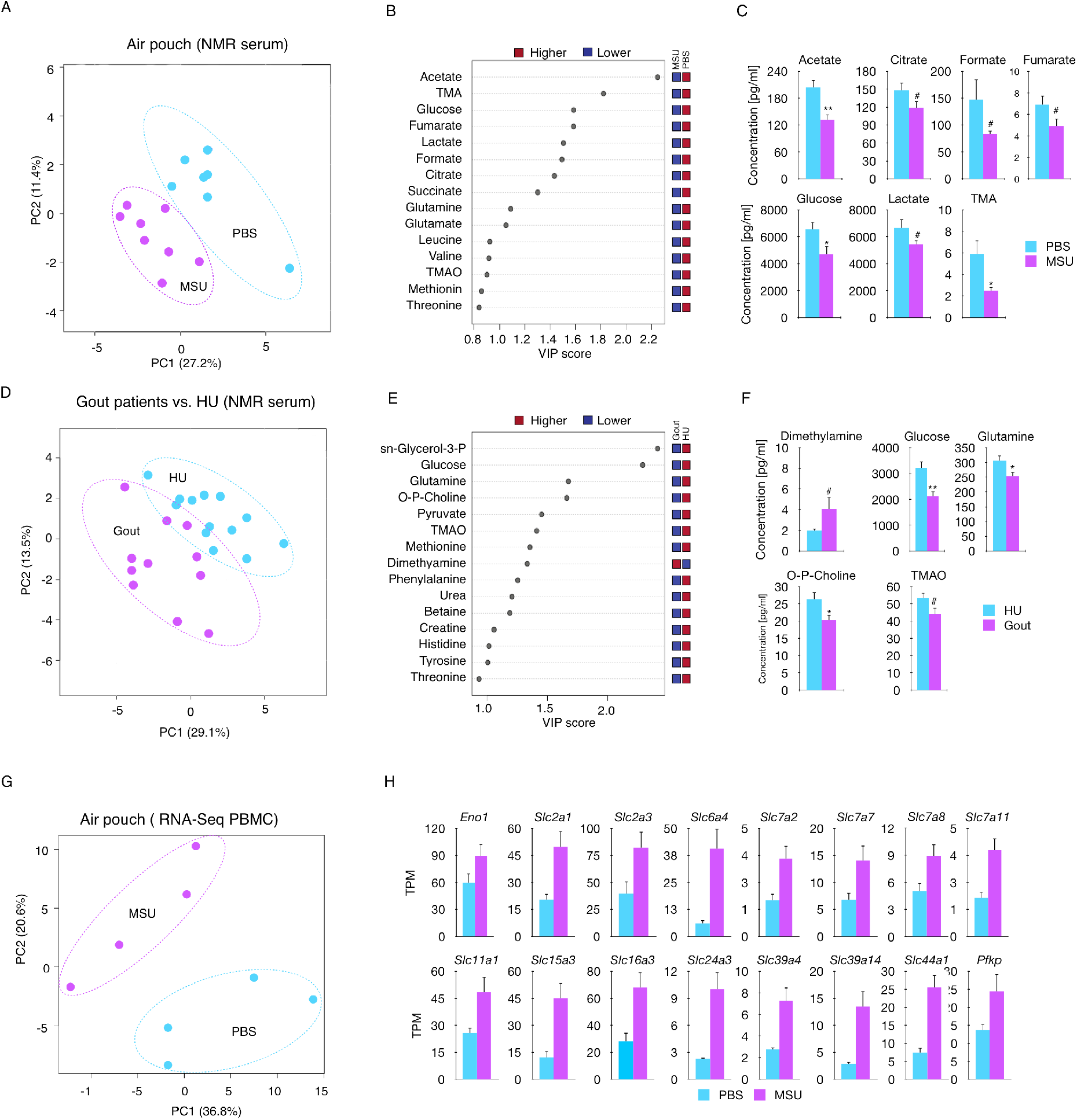
MSUc leads to systemic metabolic changes. (A) PLS-DA showing serum levels of metabolites of mice injected with PBS or MSUc in the air pouch as assessed by 1D ^1^H-NMR showing the divergence between groups (n≥7 mice/group). (B) The top 15 metabolites were ranked based on variable important projection (VIP) scores from the PLS-DA model (A). (C) Bar graphs showing reduced concentration of several metabolites including acetate, citrate, formate, fumarate, glucose, lactate and TMA in the serum of mice injected with MSUc. D) PLS-DA showing serum levels of metabolites of individuals with gout flare vs. HU as assessed by 1D ^1^H-NMR. Data shows the divergence between groups (n=11 individuals with gout and n=13 individuals with HU). (E) The top 15 metabolites were ranked based on variable important projection (VIP) scores from the PLS-DA model (A). (F) Bar graphs show reduced concentration of several metabolites including glucose, glutamine, phosphocholine and TMA and increased dimethylamine in the serum of individuals with gout. (G) PCA of the transcriptome of PBMC isolated from mice injected with PBS or MSUc in the air pouch showing the divergence between groups (n=4 mice/condition). (H) TPM of several genes involved in metabolic pathways in PBMC from mice injected with MSUc in the air pouch (P<0.05). (#=P<0.10;*=P<0.05; **=P<0.01).

Since our data indicate systemic metabolic changes induced by MSUc, we next investigated whether PBMC would have an altered transcriptional program when injected with MSUc in the air pouch. Indeed, MSUc injection in the air pouch altered the transcriptomic profile of PBMC **(Figure 3G)** including up-regulation of several genes coding for *Slc* transporters **(Figure 3H)** and inflammatory cytokines and adhesion molecules **(Supplementary Figure 3C)**. As expected, these changes were accompanied by increased concentration of cytokines CXCL1, CCL2 and IL6 in blood and CXCL1, CXCL2, CCL2 and IL6 in the air pouch lavage of mice injected with MSUc in the air pouch **(Supplementary Figure 3D,E, respectively)**. Our data indicate that MSUc triggers a metabolic program in macrophages that is also observed in animal models of gout and in serum from patients with acute gout flare. In addition, local acute gouty inflammation leads to increased cytokines levels and metabolic changes both locally and systemically, that is accompanied by an altered metabolic and inflammatory profile of PBMC.

### MSUc de-regulate a transcription factor network featured by increased AP-1

To decipher transcriptional mechanisms by which MSUc induces an inflammatory and metabolic program of gene expression, we performed *in silico* promoter (−2000 to +500 bp from the transcription start site) analysis of genes up-regulated in MSUc or LPS using HOMER. The promoters of genes up-regulated by MSUc were enriched in motifs for AP-1 members in both BMDM and mDM. Other transcription factors up-regulated by MSUc were circadian clock proteins, MYC, NFILs and the MITF/TFE family **(Figure 4A upper panel)** but not motifs for IRF, which were only displayed in the promoter of genes up-regulated by LPS **(Figure 4A lower panel)**. Interestingly, macrophages treated with MSUc displayed increased expression of several AP-1 members including *JUN, JUNB, JUND, ATF3, FOSB, FOSL1* and *FOSL2* mRNA and JUN, p-JUN^Ser63^, ATF3, FOSB, FOSL1, FOSL2 protein levels but not IRFs, which were only up-regulated in macrophages treated with LPS **(Figure 4B-D and Supplementary Figure 4)**. Some of them, including JUN, pJUN^Ser63^ and FOSL1 displayed higher up-regulation in MSUc than LPS.

**Figure 4.**
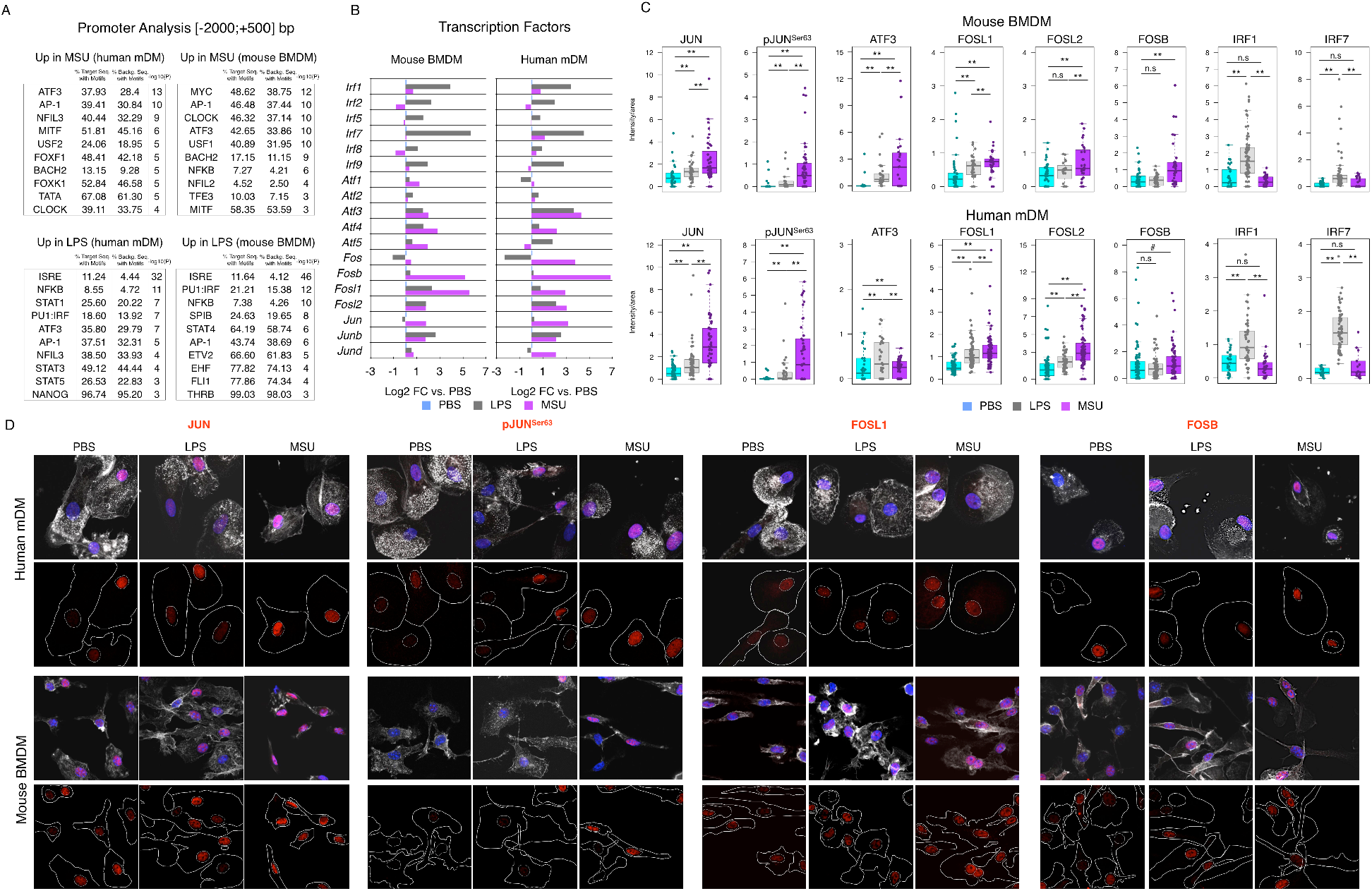
MSUc leads to activation of AP-1 but not IRFs. (A) Motif analysis using HOMER of the promoter [-2000;+500 bp, TSS] of genes up-regulated by LPS or MSUc in mDM or BMDM showing enrichment in motifs for AP-1, MYC, MITF, NRF2 and nuclear receptors –but not IRFs-in the promoters of genes up-regulated by MSUc. (B) Bar plots showing up-regulation of several AP-1 members –but not IRFs-in macrophages treated with MSUc assessed by RNA-Seq. Note the higher up-regulation of AP-1 in MSUc compared to LPS except for ATF3 in mDM. (C) Quantification of nuclear expression of JUN, pJUN^Ser63^, ATF3, FOSL1, FOSL2, FOSB, IRF1 and IRF7 in macrophages treated with MSUc or LPS showing up-regulation of AP-1 family members –but not IRF1 or IRF7-in macrophages treated with MSUc. (D) Representative images of protein analysis by IF showing up-regulation of JUN, pJUN^Ser63^, FOSL1 and FOSB in macrophages treated with MSUc or LPS used to generate plots in panel C. Note the higher up-regulation of AP-1 in MSUc compared to LPS. (#=P<0.10;*=P<0.05; **=P<0.01).

### Activation of JNK is required for the inflammatory and metabolic changes induced by MSUc

The over-representation of AP-1 motifs in the promoters of up-regulated genes by MSUc and the increased expression of several AP-1 members including JUN and pJUN^Ser63^ suggest a role of JNK in the transcriptomic program activated by MSUc. MSUc induces the up-regulation of p-JNK that is accompanied by sustained nuclear shuttling of p-JNK **(Figure 5A** and **Supplementary Figure 5A,B)** suggesting a distinct activation of JNK in MSUc vs. LPS. In line with this, treatment of BMDM with SP600125, a JNK inhibitor (JNKi), ameliorated the gene up-regulation in BMDM treated with LPS but to a larger extent in BMDM treated with MSUc (P=e^-49^ for MSUc and P=e^-34^ for LPS) (**Figure 5B)**. Whereas in LPS, JNKi ameliorated to a greater extent genes only up-regulated in LPS vs. those up-regulated in LPS and MSUc (P=e^-26^ vs. P=e^-11^, respectively), similar reduction was observed for MSUc unique up-regulated genes vs. genes similarly up-regulated by MSUc and LPS (P=e^-28^ vs. P=e^-23^, respectively)**(Figure 5C)**. Moreover, a distinct distribution of changes LPS+JNKi differed to that observed in MSUc+JNKi vs.MSUc compared to LPS+JNKi vs. LPS (**Supplementary Figure 5C)**. The promoter of genes downregulated in LPS+JNKi vs. LPS were enriched in putative binding sites for IRFs, NFKB and AP-1. Those downregulated in MSUc+JNKi vs.MSUc were enriched in AP-1, MYC, NRF2 and circadian clock proteins **(Supplementary Figure 5D)**. Whereas gene sets recovered in LPS or both LPS and MSUc belonged to inflammatory signaling pathways, those uniquely ameliorated by MSUc were related to metabolism including metabolism of carbohydrates, glycosaminoglycan, transport of small molecules and Slc-mediated transmembrane transport (**Figure 5D** and **Supplementary Figure 5E)**. Of note, treatment with JNKi only rescued partially the increased expression of metabolic genes suggesting additional mechanisms involved **(Figure 5E)**. Interestingly, several inflammatory genes including *Ccl3, Ccl9, Cxcl1, Cxcl2* and *Tnfa* that were up-regulated in LPS vs. MSUc showed higher downregulation in MSUc+JNKi vs. LPS+JNKi. Several metabolic genes including *Slc1a4, Slc2a1* and *Slc7a11* that were up-regulated to a similar extent in LPS and MSUc only showed downregulation in MSUc+JNKi. Some AP-1 members such as *Jun, Jund, Fosl1, Atf4* and *Atf5* and the stress response gene *Nqo1* were up-regulated in MSUc vs. LPS also showed higher downregulation in MSUc+JNKi vs. LPS+JNKi **(Figure 5E)**. The ameliorated up-regulation in MSUc+JNKi vs.MSUc was also demonstrated for *SLC2A1, SLC1A5, CXCL1, IL6, HK2, ENO2, SLC38A2* in mDM **(Figure 5F)**. These the mRNA changes were accompanied by reduced protein expression of JUN, JUN^Ser63^, FOSL1, FOSB and SLC2A1 in mDM **(Figure 6A,B)** and additionally PFKBP3 in BMDM (**Supplementary Figure 5F,G)** treated with MSUc vs. MSUc+JNKi as assessed by immunofluorescence and ELISA for CCL2, CXCL1, CXCL2 and TNF **(Figure 6C)**. Of note, whereas JNKi reduced partially the increased secretion of cytokines by LPS, JNKi reduced to almost basal levels the secretion of cytokines by MSUc **(Figure 6C)**. In addition, JNK inhibition also reduced the up-regulation of some of the metabolic changes observed after MSUc stimulation, specifically JNK inhibition ameliorated the increase of creatinine phosphate, dimethylamine, glutamate and lactate but not acetate, formate, pyruvate (P=0.26) or lysine **(Figure 6D)**.

**Figure 5.**
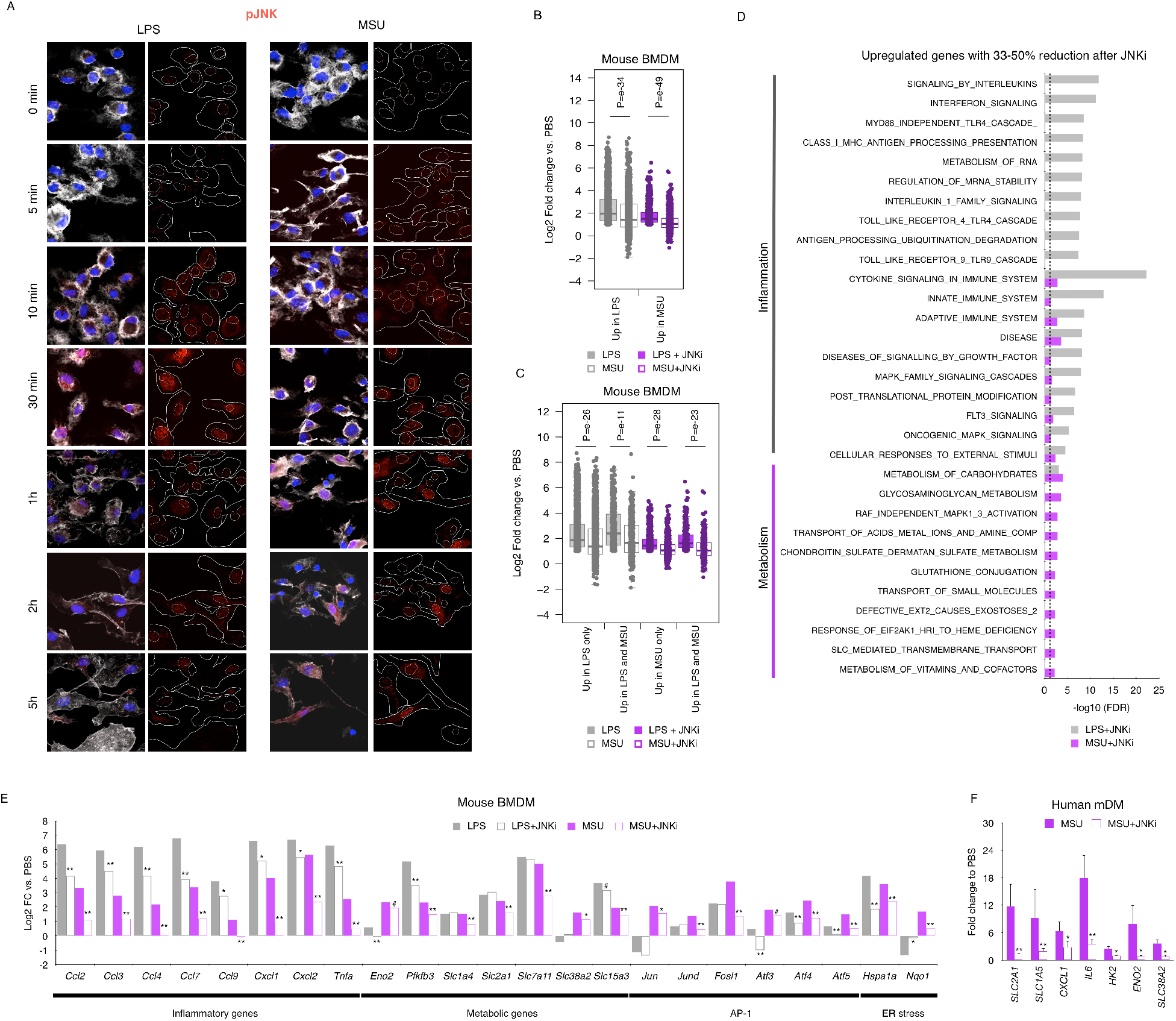
The inflammatory and metabolic program induced by MSUc is regulated through JNK. (A) Protein analysis by IF of pJNK expression in BMDM treated with LPS or MSUc at various timepoints. Data shows prolonged pJNK expression in BMDM treated with MSUc. (B,C) Box plots showing amelioration of gene expression after treatment with JNKi vs. Vehicle of genes up-regulated by LPS or MSUc (B), genes up-regulated only in LPS o rMSUc or commonly up-regulated in LPS and MSUc (C) (n≥5/group). (D) GSEA analysis using REACTOME of genes between 33%-50% reduction of expression in MSUc+JNKi vs.MSUc or LPS+JNKi vs. LPS. Data shows enrichment in inflammatory gene sets in LPS and MSUc and metabolic gene sets in MSUc. (E,F) Expression analysis by RNA-Seq (E) or RT-qPCR (F) of inflammatory and metabolic genes induced by MSUc or LPS showing reduction after treatment with JNKi in BMDM (E)(n≥5/group) or mDM (F)(n=3 donors/group). (#=P<0.10;*=P<0.05; **=P<0.01). Broken lines in D represents the cutoff for significance –log_10_(0.05)=1.30.

**Figure 6.**
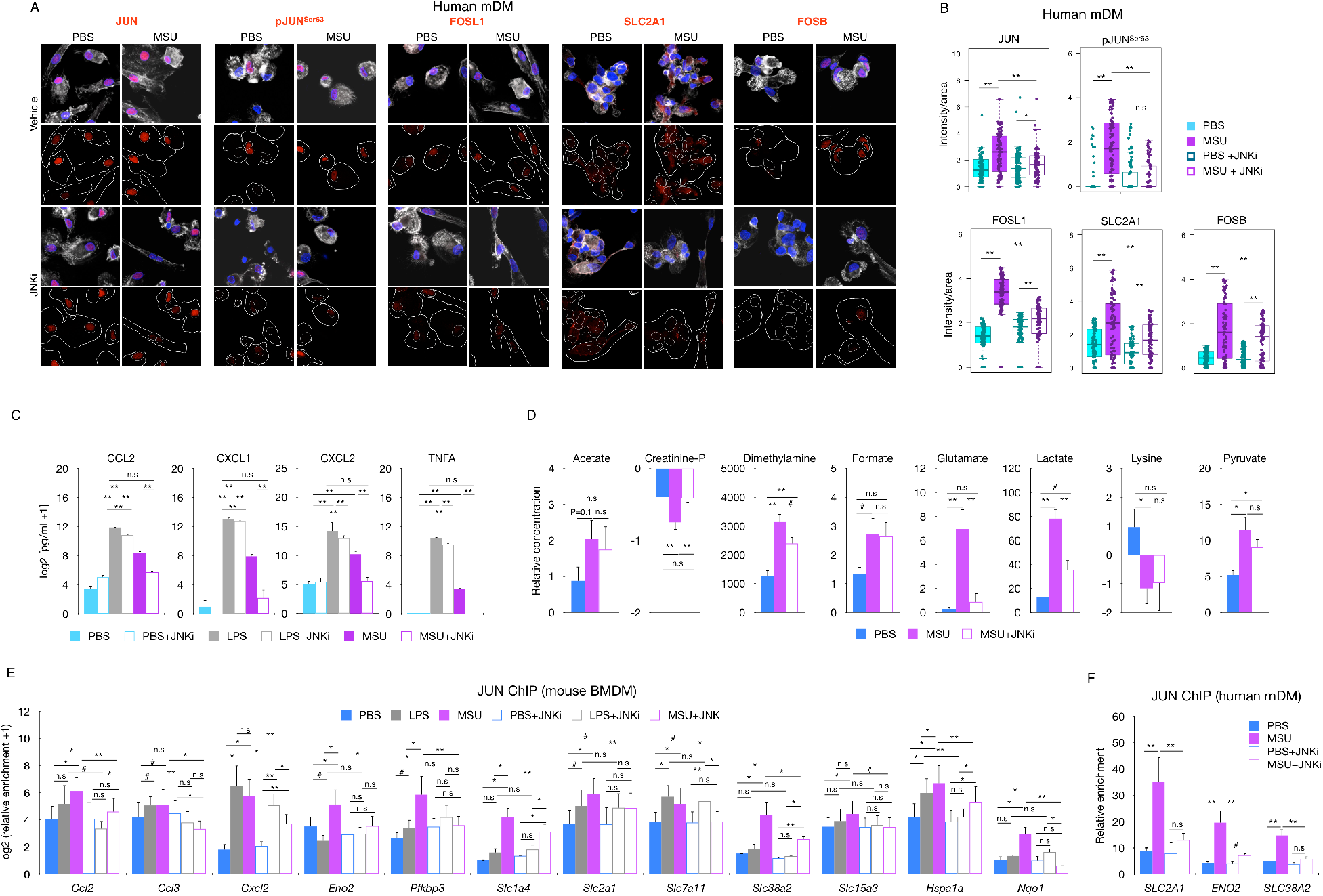
Increased JUN binding to the promoter of inflammatory and metabolic genes is regulated by JNK and is required for the activation of the inflammatory and metabolic program induced by MSUc. (A,B) Protein analysis by IF of JUN, pJUN^Ser63^, FOSL1, SLC2A1, FOSB in mDM treated with MSUc or MSUc+JNKi for 5h. Quantification of B corresponds to experiment shown in A, and shows downregulation of protein expression in MSU+JNKi vs. MSUc (n=3/condition). (C) Protein analysis by ELISA of cytokines in the supernatant of BMDM treated with LPS or MSUc w/wo JNKi showing complete reduction in MSUc+JNKi vs. MSUc and partial reduction in LPS+JNKi vs. LPS (n=3/group). (D) Analysis of metabolites by 1D ^1^H-NMR in the culture media of BMDM treated with MSUc or MSUc+JNKi showing varying degree of recovery (n=3/condition). (E,F) JUN ChIP in BMDM (E) or mDM (F) treated with LPS, MSUc, LPS+JNKi or MSUc+JNKi and qPCR over regulatory regions of genes up-regulated by LPS or MSUc. Data shows higher levels of JUN binding in macrophages treated with MSUc that is reduced upon treatment with JNKi (n=4/condition). (#=P<0.10;*=P<0.05; **=P<0.01).

Since phosphorylation by p38 is one of the major drivers of JUN phosphorylation (37), we next decided to investigate whether inhibition of p38 would have similar effect as inhibition of JNK. Interestingly, we did not observe amelioration on JUN or pJUN^Ser63^ protein levels nor reduction on up-regulated genes by MSUc in BMDM treated with MSUc and p38 inhibitor (p38i) -except for *Cxcl1*- or recovery on the up-regulation of FOSL1 or JUNB protein levels (**Supplementary Figure 6 A-D**). These data indicate that JNK –but not p38-regulates the inflammatory and metabolic program altered by MSUc.

We then investigated whether JNK dependent gene expression are due to JUN binding to the regulatory regions of up-regulated genes. We took advantage of the publicly available ATAC-Seq (38) to define regions of open chromatin in BMDM. Interestingly, even though *Ccl2, Ccl3, Cxcl2* and *Pfkbp3* are more up-regulated in LPS than in MSUc, we found similar or greater JUN enrichment in cells treated with MSUc than in LPS. In addition, genes that were only up-regulated in MSUc displayed higher JUN occupancy such as *Slc38a2* and *Nqo1*. All the genes with reduced expression in MSUc+JNKi vs. MSUc or LPS+JNKi vs. LPS showed reduced JUN recruitment to their promoter regions after JNKi including *Slc1a4, Slc2a1, Slc7a11, Slc38a2, Slc15a3* in a higher degree in MSUc+JNKi vs. MSUc. Genes that were not affected by treatment with JNKi such as *Slc1a4, Slc2a1* or *Slc7a11* in LPS+JNKi vs. LPS did not display reduced JUN recruitment after treatment with JNKi **(Figure 6E)**. Some of these findings were recapitulated in mDM treated with JNKi including reduced JUN recruitment to *SLC2A1, ENO2* and *SLC38A2* promoters **(Figure 6F)**. Altogether, our data suggest that the inhibition of up-regulated gene transcripts after MSUc by JNKi –but not p38i-is regulated through JUN recruitment to their promoter regions to a larger extent in genes up-regulated by MSUc than by LPS.

### Reduction of oxidative stress or blockage of inflammasome does not affect the activation of inflammatory or metabolic genes by MSUc

The interaction of MSUc with the plasma membrane of the cell promotes a cellular response that includes the production and release of reactive oxygen species and subsequent endoplasmic reticulum (ER) stress and oxidative stress (39, 40, 41. In line with this, we found that ER stress, activated unfolded protein response (UPR), and NRF2 target genes were up-regulated by MSUc (including *Mafg, Atf4, Hsp70* and *Soat2* mRNA and protein levels), and were also reduced by treatment with JNKi (**Figure 7A-C**). However, treatment of BMDM with the antioxidant mixture butylated hydroxyanisole (42) reduced only partially the up-regulation of JUN or pJUN^Ser63^ protein levels (21% reduction in MSU+BHA vs. MSU, P<0.10) and did not alter the over-expression of inflammatory and metabolic genes by MSUc (**Figure 7D-F**), suggesting that JUN activation by oxidative stress or ER stress is not the principal etiology of the changes induced by MSUc. Moreover, despite the role of NLRP3 inflammasome in gout being well described (18, 21), we found that treatment of BMDM with MCC950, a NLRP3 inflammasome inhibitor, did not reduce the expression of JUN, pJUN^Ser63^ and FOSL1 protein levels nor affected the up-regulation of gene expression by MSUc (**Figure 7G**), suggesting that activation of the NLRP3 inflammasome is not required for up-regulation of JUN and transcriptomic changes induced by MSU.

**Figure 7.**
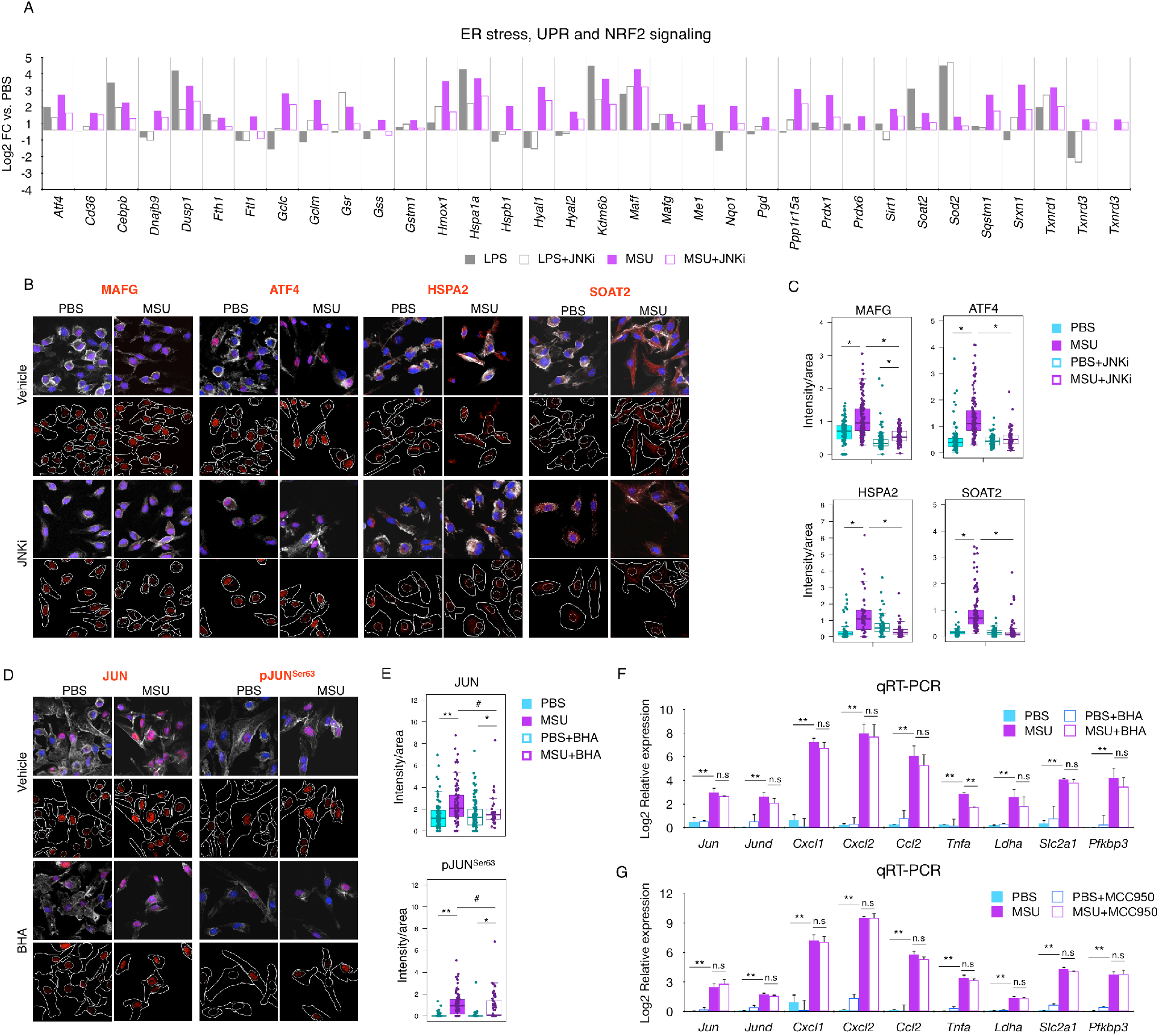
Up-regulation of the ER stress, UPR and NRF2 transcriptional program by MSUc is reduced after treatment with JNKi. (A) Expression analysis by RNA-Seq of genes involved in ER stress, UPR or NRF2 signaling after treatment with LPS, MSUc, LPS+JNKi or MSUc+JNKi showing greater down-regulation in MSUc+JNKi vs. MSUc compared to LPS+JNKi vs. LPS. (B) Protein analysis by IF of MAFG, ATF4, HSPA2 and SOAT2 in BMDM treated with MSUc or MSUc+JNKi showing down-regulation in MSUc+JNKi. (C) Quantification of experiment shown in (B). (D) Protein analysis by IF of JUN and pJUN^Ser63^ in BMDM treated with MSUc or MSUc+BHA showing partial downregulation in MSU+BHA. (E) Quantification of experiment shown in (D). (F,G) RT-qPCR analysis of inflammatory and metabolic genes in BMDM treated with MSUc or MSUc+BHA (F) MSUc or MSUc+MCC950 (G) showing minor downregulation of MSUc+BHA or MSUc+MCC950 vs. MSUc. (#=P<0.10;*=P<0.05; **=P<0.01).

### JNK and SLC2A1 transport inhibition ameliorated inflammation in animal models of acute gout

The previous results led us to investigate whether activation of JNK was required for the phenotype induced by MSUc *in vivo*. In addition, since glucose metabolism was one of the top metabolic pathways deregulated by MSUc in both BMDM and mDM, we also tested whether the inhibition of the glucose transporter SLC2A1 improved animal models of gouty inflammation. We found increased expression of pJNK, SLC2A1, LDHA, ENO2 and iNOS in macrophages in the subcutaneous air pouch lining of mice injected with MSUc **(Figure 8A,B, and Supplementary Figure 7A)**. Injection of MSUc in the air pouch lining led to a recruitment of leukocytes that was significantly lowered after treatment with JNKi (**Figure 8 C-E)** or BAY-767 **(Figure 8F-H)**. Similarly, intra-articular injection of MSUc led to increased infiltrates of inflammatory cells that were significantly reduced after treatment with JNKi **(Figure 8I,J)** or BAY-767 **(Figure 8K,L)**. Of note, we found that treatment of BMDM with BAY-876, did not reduce the expression of JUN and pJUN^Ser63^ protein levels nor affected the upregulation of gene expression by MSUc (**Supplementary Figure 7B-D**), suggesting that SLC2A1 is a critical downstream target gene of MSUc but glycolysis activation is not responsible for upregulation of JUN and transcriptomic changes induced by MSU.

**Figure 8.**
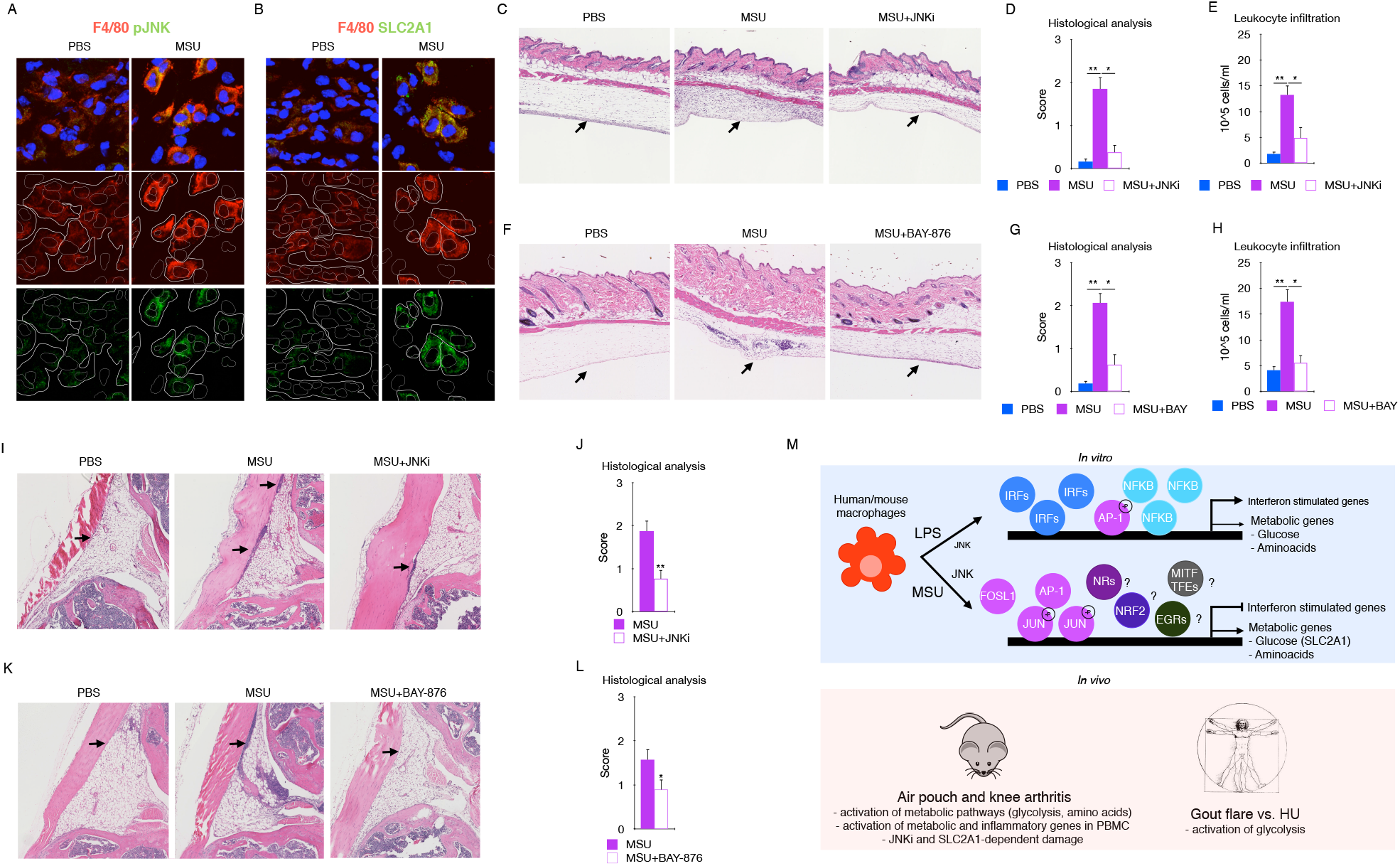
Signaling by JNK and SLC2A1 is required for the MSUc-induced damage *in vivo*. (A,B) Protein analysis by IF showing expression of SLC2A1 (A) and pJNK (B) in macrophages in the subcutaneous cavity of the air pouch when injected with MSUc (n=2/group). (C) Histological analysis by HE showing the recruitment of inflammatory cells (arrows) induced by MSUc in the air pouch is reduced upon treatment with JNKi (n≥4/group). (D,E) Pathological assessment of inflammatory cell infiltrates of the air pouch (D) and neutrophil count of the air pouch lavage (E) showing reduction in MSUc+JNKi vs. MSUc (n≥4/group). (F) Histological analysis by HE showing the recruitment of inflammatory cells (arrows) induced by MSUc in the air pouch is reduced upon treatment with BAY-876 (n≥5/group). (G,H) Pathological assessment of inflammatory cell infiltrates of the air pouch (G) and neutrophil count of the air pouch lavage (H) showing reduction in MSUc+BAY-876 vs. MSUc (n≥10/group). (I,J) Histological analysis by H&E showing reduction in the recruitment of inflammatory cells (arrow) in the synovial cavity of mice injected with MSUc+JNKi vs. MSUc (n≥10/group). (K,L) Histological analysis by H&E showing reduction in the recruitment of inflammatory cells (arrow) in the synovium of mice injected with MSUc+BAY-876 vs. MSUc (n≥7/group). (M) Diagram showing that MSUc induces a unique transcriptional program with up-activation of inflammatory –but not interferon stimulated genes- and metabolic genes, which is regulated through up-regulation of AP-1 via JNK but not IRFs. (#=P<0.10;*=P<0.05; **=P<0.01).

Our data indicate that MSUc activates a distinct inflammatory and metabolic program in macrophages featured by increased glycolysis and amino acid metabolism, which is regulated by JNKi through binding of JUN to the promoter of target genes. In addition, although oxidative stress, inflammasome and SLC2A1 are critical signatures induced by MSUc, they are not responsible for the transcriptomic changes induced by MSU. Finally, our data has clinical relevance since serum of individuals with acute gout flare vs. HU showed altered metabolic profile **(Figure 8M, model)**.

## DISCUSSION

The transcriptional mechanisms underlying macrophage response to MSUc are largely unknown. Here, the investigation of genome-wide transcriptomic and bioinformatic analysis of the promoter of deregulated genes in macrophages revealed that MSUc activates a distinct inflammatory and metabolic program that is regulated through transcriptional activation, protein phosphorylation, and recruitment of JUN to the regulatory regions of inflammatory and metabolic genes, and this is regulated via JNK activity.

MSUc activates the inflammatory program of macrophages. However, priming cells either with LPS or other TLRs agonists hampers very much the understanding of the response to MSUc by macrophages. Our work demonstrates up-regulation of inflammatory and metabolic genes in both mDM and murine BMDM by MSUc alone. Several metabolic pathways were up-regulated after MSUc stimulation including carbohydrates metabolism, lipid metabolism and also nutrient, including amino acid, transport by the solute carrier (SLC) transporter family. The metabolic changes featured by SLC2A1-mediated glucose uptake yield crucial insights into how MSUc gouty inflammation exerts its pathogenic effect. Whereas some of those metabolic genes are similarly de-regulated in macrophages treated with LPS mRNA, these macrophages do not display large deregulation on metabolite production (except for increased concentration of lactate), suggesting that alternative mechanism to mRNA expression –or different timing-govern the control of metabolic pathways by LPS including protein stability and membrane localization. Additional experiments using fine resolution confocal microscopy and immunogold staining followed by electron microscopy would be helpful to address these points.

Amino acid pathways such as glycine, serine and threonine, and phenylamine pathway, and amino acid related SLC transporters were more obviously deregulated after MSUc than LPS stimulation, both in gene expression and metabolite levels. Prior studies have highlighted the important role of amino acids in macrophage phenotype and function. Amino acids including glutamine, arginine or glycine were shown to control macrophage polarization and used differently in different macrophage subtypes. Some of these metabolic changes controls macrophage polarization by regulating epigenetically the transcriptional activity of key genes (43-45). Whether MSUc activates a unique macrophage phenotype with a particular metabolic rewiring that helps to inflammation self-limitation and gout resolution or whether targeting other SLC transporters play a key role in gout pathogenesis need further studies. In addition, stable isotope tracing metabolomics would be critical to understand the amino acid metabolic pathway activity, via the relative production and consumption of the labeled metabolites.

Interestingly, local acute inflammation by MSUc altered not only the sera metabolic profile (46, 47) but it also increased expression of metabolic and inflammatory genes in PBMC suggesting a systemic effect. Our data from mouse has clinical implications since we demonstrated that individuals with gout display lower levels of blood glucose levels when compared to those with HU but not gout. Moreover, prior studies have shown 75 transcription factor motifs associated with hypomethylated differential methylation loci regions in gout vs. healthy controls (48) including those that belong to MEF2C, NFATC2, JUN and FOS, HIF1A, MYC-MAX and circadian clock regulators CLOCK and BHLHLE40. Interestingly, we find upregulation of *Fosb, Fosl1, Fosl2, Hif1a, Max* and *Bhlhe40* –but not *Mef2c* that was downregulated or *Nfatc2* that was unaffected-in PBMC from mice injected with MSUc indicating that similar signaling pathways might be involved in the activation of PBMC in patients with gout during flares. Interestingly, analysis of promoter of up-regulated genes in PBMC from mice injected with MSUc in the air pouch show slight enrichment for AP-1 motifs suggesting that, although of importance, AP-1 might not be the major driver of gene expression in PBMC (data not shown). Dysregulated trained immunity can be critical in the context of chronic inflammatory disorder. And is known that mechanisms driving trained immunity include profound changes in intracellular metabolism, which are associated with epigenetic reprogramming (5, 49). Whether or not MSUc is able to build a long-term pro-inflammatory phenotype after primary stimulation in monocytes or is able of reprogramming of myeloid progenitor cells in the bone marrow is unknown at this point.

Previous studies have shown the inflammatory program triggered by MSUc (18, 50, 51), and recently a work by Renaudin and colleagues (22) found that MSUc favored glycolytic activity of primed THP-1 cells. However, Renaudin’s work is limiting in identifying the mechanism involved in the MSUc effects in primary mouse and human macrophages. We provide evidence that AP-1 family signaling is required for activation of transcriptional program by MSU. Enrichment of AP-1 motifs in the promoter of up-regulated genes by MSUc was accompanied by up-regulation of the majority of AP-1 members at mRNA and protein level, including phosphorylation of JUN at the Ser^63^ residue. Activation of AP-1 activity is modulated through its dimer composition (52, 53), which is determined by the regulation of the synthesis and stability of respective mRNA or through the regulation of protein stability (for example via phosphorylation). Phosphorylation of AP-1 members modulates the transcriptional activity of AP-1 dimers (54) in a process that is mostly mediated by the JNK and p38 MAPK pathways that once translocated to the nucleus regulates the transcriptional activity of AP-1 dimers (55). Therefore, mechanisms that activate phosphorylation and/or mRNA synthesis would change the stoichiometry of AP-1 members and would dictate their dimer composition.

We found no evidence that MSUc induces an hyperphosphorylation of JNK compared to LPS. However, the prolonged nuclear pJNK cytosolic and nuclear shuttling in BMDM exposed to MSUc suggests that increased pJUN^Ser63^ could be due both to the unexpected increased expression of JUN mRNA, and/or increased shuttling of pJNK to the nucleus where JUN interacts with other AP-1 members. Of interest, steroidal and non-steroidal anti-inflammatory drugs that are used to treat acute flares of gout are known to abrogate JNK signaling (56, 57). Our finding that treatment with p38i has no effect in reducing JUN/pJUN^Ser63^ nor to ameliorate the transcriptomic effects induced by MSUc, suggest that p38 is not the main mechanism of JUN phosphorylation by MSUc. Regardless of the mechanism, phosphorylation of JUN on Ser 63/73, for example, potentiates its transcriptional activity by heterodimerization with FOS (58, 59). Interestingly, we find that *FOS* is up-regulated by MSUc but not by LPS (Log2 fold change -1.08 vs. 0.5 in BMDM or -2.31 vs. 3.77 in mDM after LPS or MSUc, respectively) and *FOSB* is the AP-1 member with largest up-regulation in MSUc compared to LPS (Log2 fold change 0.44 vs. 5.19 in BMDM or 0.196 vs. 6.85 in mDM after LPS or MSUc, respectively). Transactivation of JUN is also potentiated by ATF2 through the binding of Rb or E1A to the ATF2-JUN dimer (37, 60, 61). However, we find no up-regulation of *ATF2* in macrophages stimulated with MSUc suggesting that this might not be the major cause of increased JUN activity.

Thus, our data point to a strong transcriptional activation of AP-1 genes by MSUc. The expression of several AP-1 components is under positive and negative AP-1 autoregulation and this feedback allows a finely tuned regulation of AP-1 members activity over time (62-65). In pancreatic epithelial cells, for instance, *Jun* mRNA expression is regulated thorough the cooperative binding of the nuclear receptor NR5A2 and JUN itself (66) suggesting that nuclear receptors might be playing a role in *Jun* up-regulation by MSUc.

Our results using JUN ChIP demonstrate that the recruitment of JUN to the promoter of up-regulated genes is greater in MSUc than LPS regardless of the degree of up-regulation of the gene transcript. Interestingly, treatment with JNKi has greater effect over the up-regulated genes induced by MSUc than by LPS. And, of those induced by LPS, JNKi has a greater effect in LPS-unique up-regulated genes compared to those also up-regulated by MSUc suggesting that even when a gene is deregulated both by LPS and MSUc, JNK has greater importance for its regulation in MSUc than LPS. Our data provide evidence that treatment with JNKi could be therapeutically beneficial to overcome the damage associated with gouty inflammation.

Dysregulation of JNK pathway is associated with a wide range of immune and neurological disorders and cancer, and there is an active research to develop better small molecules to inhibit JNK without generating major side effects (42, 67-72). The motif analysis of the promoters of genes that are downregulated with JNKi vs. vehicle suggest that whereas the effect of JNKi on LPS-activated genes might be secondary to activation of IRFs; the effect of JNKi in MSUc-activated genes is mainly driven through modulation of AP-1 activity in cooperation with other transcription factors such as NRF2, MITF and circadian clock regulators.

Although a critical initial step is provided by the up-regulation of JUN and its phosphorylation by JNK, it is expected that other transcription factors take part in the transcriptional program induced by MSUc. Interestingly, we found that MSUc induced the up-regulation of the EGR1, EGR2 and EGR3 (data not shown), MYC and transcription factors of the circadian clock family such as BHLHE40, PER1, NR1D1, CRY1. MYC regulates several metabolic and inflammatory genes in macrophages (30, 73, 74). Interestingly, preliminary ChIP-Seq experiment using antibodies against H3K27ac showed enriched motifs for AP-1, circadian clock, EGRs, NRF2 and MYC in the enhancers activated by MSUc suggesting a role in the regulation of enhancer activity (data not shown). The presence of collaborative binding between other TF might explain the partial rescue of the metabolic program induced by MSUc. Interestingly, preliminary experiment also show that damage induced by MSUc injection in the joint is significantly reduced upon injection of MYC inhibitors (data not shown), indicating that MYC activity is required *in vivo* for the damage induced by MSUc. Experiments using macrophage-specific knock-out mouse models would be helpful to address which transcription factor network is required for the inflammatory and metabolic alterations induced by MSUc *in vitro* and *in vivo*.

Despite the role of NLRP3 inflammasome in gout disease is well described we found no evidence that the inflammasome is required for the up-regulation of inflammatory and metabolic genes by MSUc. We also found downregulation of *Tlr3, Tlr8, Tlr9* mRNA in BMDM treated with MSUc and downregulation of *TLR4, TLR5* and *TLR6* mRNA in mDM treated with MSUc, that together with the significant differences with LPS program –including the lack of activation of IFN program by MSU-, suggest that activation by TLRs is not the mayor driver of the phenotype observed in macrophages.

Finally, gout attacks are self-limiting suggesting the certain regulatory mechanisms are present to modify the acute inflammatory response. Preliminary data showed that MSUc shares a larger fraction of inflammatory genes with LPS at shorter time-points suggesting a transcriptional repression of the inflammatory program at later time-points. Genome-wide analysis of the transcriptomic effects of MSUc and LPS at different time-points would help to address these points. Our data calls for an alternative phenotype of macrophages exposed to MSUc that is not similar to that phenotype induced by LPS or IL4 (data not shown). Isolation of macrophages from the synovial cavity of patients with gout would be helpful to phenotypically characterize the effect of MSUc in resident macrophages.

## MATERIAL AND METHODS

### Reagents

PBS, DMEM (w/5% glutamine, 10% FBS, 5% penicillin), trypsin 0.25%, LPS (100 ng/mL), MSUc were prepared as described (23) suspended at 25 mg/mL in sterile, endotoxin-free phosphate buffered saline (PBS) and verified to be free of detectable lipopolysaccharide contamination by Limulus lysate assay (Lonza, Walkersville, Maryland, USA), MCSF (10 ng/mL, Peprotech), GLUT1 inhibitor BAY-876 (20nM, Selleckchem, #S8452), p38 MAPK inhibitor SB203580 (3 uM, Tocris, #1202), JNK inhibitor SP600125 (20uM, Tocris, #1496), beta-hydroxybutyric acid (BHA; 10uM, Sigma, #B1253), NLRP3 inflammasome inhibitor MCC950 (1uM, Selleckchem, #S7809).

### Human monocyte derived macrophages (mDM)

Human peripheral blood mononuclear cells (PBMC) were isolated from whole blood by Ficoll density gradient using Ficoll Plaque Premium (Sigma GE Healthcare, #17-544-02) or BD Vacutainer CPT Tubes (BD, 362753) as described somewhere else. PBMC were washed twice with HBSS (Gibco ThermoFisher, #14175095) containing 2% BSA, 1mm EDTA. Monocytes were obtained by negative selection using a kit (StemCell, #19359). Monocytes were incubated with RPMI 1640 (Sigma Aldrich, #10-040) medium containing 10% de-complemented FBS (OmegaScientific, #FB-02), 1mM Sodium Pyruvate (Gibco Thermofisher), Penicillin-Streptomycin (15140122, 1000U/ml) and 25ng/ml-50ng/ml of recombinant human M-CSF (StemCell #78057-2) for six days at 37°C and 5% CO_2_. Fresh media was added 48h and four days after seeding. The mDM were incubated for five hours with 100 ng/ml of Kdo2 lipid KLA or 250 μg/mL of MSUc in RPMI 1640 medium containing 10% de-complemented FBS, 1mM Sodium Pyruvate, Penicillin-Streptomycin and 25ng-50ng/ml of recombinant human M-CSF. 20 μM of JNK inhibitor SP600125 (Tocris, #1496) was added to the media one hour prior treatment with MSUc.

### Bone-marrow derived macrophages (BMDM)

Flushed bone marrow from both femurs and tibias of C57BL/6 mice were cultured in DMEM medium supplemented with 10% de-complemented FBS, 100 units/ml penicillin and 100 ug/ml streptomycin, 2mM L-glutamine, and 20% of L929 conditioned media for for six days at 37°C and 5% CO_2_. On day six, BMDM were trypsinized and plated in DMEM medium supplemented with 10% de-complemented FBS, 2mM L-glutamine, 100 units/ml penicillin and 100 ug/ml streptomycin, and 10 ng/mL recombinant murine M-CSF (Peprotech, #315-02) for one day prior being stimulated for five hours with 100ng/mL of LPS or 250 μg/mL of MSUc. 20 μM of JNK inhibitor SP600125 (Tocris, #1496), 10 μM of BHA (Sigma, B1253), 20nM of BAY-876 (Selleckchem,#S8452), 3 μM of p38 MAPK inhibitor SB203580 (Tocris, #1202) or NLRP3 inflammasome inhibitor MCC950 (1uM, Selleckchem, #S7809) were added to the media one hour prior to stimulation with LPS or MSUc.

### Subjects

Patients meeting the 2015 ACR/EULAR gout classification criteria with hyperuricemia were recruited. Consecutive patients were recruited from the Rheumatology Outpatient Clinic. The study was approved by the VA Institutional Board Review and patients signed an informed consent. Patients with not known diabetes or dyslipidemia were included for NMR analysis. Non-fasting blood samples were collected in the clinic by research personnel into 10 ml BD Vacutainer Blood Collection Tubes containing spray-coated silica and a polymer gel for serum separation. After 30 min incubation at room temperature, tubes were centrifuged for 10 min at 2000×g and sera were transferred into 1.7ml tubes and immediately frozen and stored at −80 °C until analysis.

### RNA-Seq

It was performed as described elsewhere (24, 25). Briefly, BMDM or mDM were lysed in TRIzol (ThermoFisher, #15596026). RNA isolation and DNase treatment was carried out using Direct-zol RNA MicroPrep kit (Zymoresearch, #11-33MB). 500ng-1 μg total RNA was enriched in poly-A tailed RNA transcripts by double incubation with Oligo d(T) Magnetic Beads (NEB, S1419S) and fragmented for 9 min at 94°C in 2X Superscript III first-strand buffer containing 10mM DTT (Invitrogen, #P2325). The 10 μl of fragmented RNA was added to 0.5 μl of Random primers (Invitrogen, #48190011), 0.5 μl of Oligo d(T) primer (Invitrogen, #18418020), 0.5 μl of SUPERase inhibitor (Ambion, #AM2696), 1 μl of 10mM dNTPs and incubated at 50°C for three minutes. Then, 5.8 μl of water, 1 μl of 10mM DTT, 0.1 μl of 2 μg/μl Actinomycin D (Sigma, #A1410), 0.2 μl of 1% Tween-20 (Sigma) and 0.5 μl of SuperScript III (Invitrogen, #ThermoFisher 18080044) was added to the mix. Reverse-transcription (RT) reaction was performed at 25°C for 10 min followed by 50°C for 50 min. RT product was purified with RNAClean XP (Beckman Coulter, #A63987) and eluted in 10 μl in 0.01% Tween-20. The RNA-cDNA complex was then added to 1.5 μl of 10X Blue Buffer (Enzymatics, #B0110-L), 1.1 μl of dUTP mix (10 mM dATP, dCTP, dGTP and 20 mM dUTP), 0.2 μl of 5U/μl RNAseH (Enzymatics, #Y9220L), 1 μl of 10U/μl DNA polymerase I (Enzymatics, #P7050L), 0.15 μl of 1% Tween-20 and 1.05 μl of nuclease free water; and incubated at 16°C for 2.5h or overnight. The resulting dsDNA product was purified using 28 μl of SpeedbBead Magnetic Carboxylate (GE Healthcare, #651521050 50250) diluted in 20%PEG8000:2.5M NaCl to a final 13% PEG concentration, washed twice with 80% etOH, air dry and eluted in 40 μl of 0.05% Tween-20. The purified 40 μl of dsDNA was end-repaired by blunting followed by A-tailing and adapter ligation as described elsewhere (26) using BIOO Barcodes (BIOO Scientific, #514104), IDT TruSeq Unique Dual Indexes or Kapa Unique Dual-Indexed Adapters using 15 μl Rapid Ligation Buffer (Enzymatics, #L603-LC-L), 0.33 μl 1% Tween-20 and 0.5 μl T4 DNA ligase HC (Enzymatics, #L6030-HC-L). Libraries were amplified by PCR for 11-15 cycles using Solexa IGA and Solexa IGB primers (AATGATACGGCGACCACCGA and CAAGCAGAAGACGGCATACGA, respectively), purified using 1 μl of SpeedbBead Magnetic Carboxylate in 15.2 μl of 20%PEG8000:2.5M NaCl, washed with 80% etOH and eluted in 0.05% Tween-20. Eluted libraries were quantified using a Qubit dsDNA HS Assay Kit and sequenced on a NextSeq 500 or Hi-Seq 4000 (Illumina, San Diego, California). A list of the RNA-Seq samples can be found in Supplementary Table 5.

### Analysis of RNA-Seq

FASTQ sequencing files were mapped to the mm10 or hg38 reference genomes for mouse samples or human samples, respectively using STAR with default parameters. Biological replicates were used in all experiments. Quantification of transcripts was performed using rnaQuan.R with parameter -l 200 and analyzeRepeats.pl (HOMER) with parameters -condenseGenes -count exons -noadj. Principal Component Analysis (PCA) was obtained based on the Transcripts Per kilobase Million (TPM) on all genes of all samples. Differential expression analysis was performed using rnaDiff.R for genes with a minimum TPM of 0.5 in at least two samples. Genes with log2 fold chance > 1 or < (−1) and FDR < 0.05 were considered as differentially expressed. Only libraries with >80% mapped reads were used for downstream analysis. A list of differentially expressed genes in human mDM and mouse BMDM is provided in Supplementary Table 1 and 2, respectively.

### Gene Set Enrichment Analysis (GSEA)

Analysis of signaling pathways regulated by differentially expressed genes was performed with the Molecular Signature Database (MSigDB) of GSEA (27-29) computing the overlaps with REACTOME, HALLMARKS or KEGG datasets as indicated in the Figure legends. A list of the gene sets of genes differentially expressed in human mDM or mouse BMDM is provided in Supplementary Tables 3 and 4, respectively.

### Promoter scanning analysis

*In silico* promoter analysis of differentially expressed genes was performed using the findMotifs.pl tool of HOMER searching for motifs of length between 8 and 10 nucleotides and from -2000 to +500 bp relative to the TSS, using 4 threads.

### Gene expression analysis by retrotranscription and quantitative PCR (RT-qPCR)

Between 500K and 2.10^6^ cells were lysed in Trizol (Invitrogen, #15596026). Total RNA was isolated using Direct-zol RNA MicroPrep kit (Zymoresearch, #11-33MB) and treated with DNaseI on column. 500 ng-1μg of RNA solution was enriched in poly-A tailed RNA by two consecutive incubations with Oligo d(T) Magnetic Beads (30S1419S) or directly used for cDNA synthesis. cDNA was prepared according to the manufacturer’s specifications, using the SuperScript III Reverse Transcriptase (ThermoFisher, #18080093) or Superscript VILO cDNA master mix (Thermofisher, #11756050). RT-qPCR analysis was performed using the KAPA SYBR Fast PCR master mix (Sigma Aldrich, #KK4605) or SYBR Green PCR Master Mix (Applied Biosystems, #43-091-55) and an ABI-StepOne 96 well plate instrument (Applied Biosystems). Expression levels were normalized to *HPRT* mRNA levels using the ΔΔ*C*t method. The sequence of the primers used can be found in Supplementary Table 5).

### Chromatin immunoprecipitation (ChIP)

JUN ChIP was performed as described somewhere else (25, 31). Briefly, BMDM or mDM were fixed with 3mM Disuccinimidyl-glutarate, DSG, (Proteochem, #C1104) in PBS for 30 min at room temperature followed by 10 min incubation with 1% formaldehyde (ThermoFisher, #28906) at room temperature. Then, fixation was quenched by adding 2.625 M of glycine to a final concentration of 125 mM. Cells were washed with 0.01%Triton-X-100:PBS, scraped and centrifuged for 10 min at 3,000 rpm at 4°C and then washed once again with 0.01%Triton-X-100:PBS, pelleted, snap frozen and stored at -80°C. For ChIP experiments, cells were thawed and permeabilized in 1 ml of ice-cold buffer containing 10mM HEPES/KOH pH7.9, 85mM KCl, 1mM EDTA, 0.2% IGEPAL CA-630 (Sigma Aldrich, #I8896), 1X protease inhibitor cocktail (Sigma, #11836145001) and 1 mM PMSF for 10 min on ice. Cells were then spun down and lysed in 130 μl of lysis buffer containing 20 mM Tris/HCl pH7.5, 1 mM EDTA, 0.5 mM EGTA, 0.1% SDS, 0.4% Sodium Deoxycholate, 1% NP-40, 0.5 mM DTT, 1x protease inhibitor cocktail and 1 mM PMSF and chromatin was sheared by sonication. In all buffers, DTT, protease inhibitor and PMSF were added freshly. Cell lysates were sonicated in a 96 microTUBE Rack (Covaris, #500282) using a Covaris E220 for 22 cycles with the following settings: time, 60 seconds; duty 5.0; PIP, 140; cycles, 200; amplitude, 0.0; velocity, 0.0. Sonicated lysates were recovered and spun at 10,000 rpm for 10 min at 4°C to remove cell debris. One percent of sonicated lysate was kept as ChIP input for analysis. Immunoprecipitation mix consisting of Protein G Dynabeads (Invitrogen, #10003D) and 0.2 μg of JUN antibody (abcam, #32127) was added to sonicated chromatin solution and incubated overnight on a rotator at 4°C. Next day, immunocomplexes were placed on a magnet and bead complexes were washed for one minute with 150 μl of cold buffers as indicated: 3 times lysis buffer, 4 times with wash buffer containing 10 mM Tris/HCl pH7.5, 250mM LiCl, 1 mM EDTA, 0.7% Na-Deoxycholate and 1% NP-40 alternative; 2 times with TET buffer containing 10 mM Tris/HCl pH 8.0, 1 mM EDTA, 0.2% Tween-20; and 1 time with IDTE buffer containing 10 mM Tris/HCl pH 8.0 and 0.1 mM EDTA. Bead complexes were resuspended in 25 μl of TT buffer containing 10 mM Tris/HCl pH 8.0, 0.05% Tween-20. All wash buffers contained 1X Protease Inhibitor cocktail except for TT buffer. Crosslinks were reversed by adding 33.5 μl of mix containing 18.4 μl nuclease free water, 4 μl 10% SDS, 3 μl 0.5 M EDTA, 1.6 μl 0.2M EGTA, 1 μl 10mg/ml Proteinase K (Biolabs, #P8107S), 1 μl 10mg/ml RNase A (ThermoFisher, #12091021), 4.5 μl 5M NaCl by incubating at 55°C for 1h followed by 75°C for 30 min. Dynabeads were removed and libraries were cleaned by adding 2 μl of SpeadBeads in 124 μl of 20% PEG 8000/1.5M NaCl, washed by adding 150 μl 80% EtOH, air dried and eluted in 15 μl of buffer containing 10 mM Tris/HCl pH 8.0 and 0.05% Tween-20. Enrichment was calculated as relative to input and to the negative region. A list of primers used for ChIP-qPCR is provided in Supplementary Table 5.

### Immunofluorescence (IF) in cells

Briefly, cells were plated in chambered slides after seven days of differentiation for murine BMDM or differentiated directly in chambered slides (Millipore, #C86024) in the case of human mDM. Cells were fixed with BD Cytofix/Cytoperm Buffer (BD, #BD554714) or 4% PFA containing 0.1% Saponin (Sigma, #47036) for 10 min at room temperature and then washed twice with HBSS containing 2% BSA and 1mM EDTA. Cells were incubated in wash/permeabilization buffer (BD, #BD554714) for one hour at 4°C or kept at 4°C until the experiment was performed. Fixed BMDM were blocked using 3% BSA, 0.1% Triton-PBS for 30 min at room temperature and then with the primary antibody overnight at 4°C. For double IF, the corresponding antibodies were added simultaneously and incubated overnight at 4°C. Next day, mDM were washed with 0.1% Triton-PBS, incubated with the appropriate fluorochrome-conjugated secondary antibody and fluorescent probes. Nuclei were counter-stained with DAPI and Phalloidin (Abcam, #176759) was used to identify cell perimeter. After washing with 0.1% Triton-PBS, slides were mounted with Prolong Gold Antifade Reagent (Life Technology, #10144). Images were taken using a Leica SP8 with light deconvolution microscope or Leica TCS SPE microscope. For antibodies raised in mouse signal was pseudocolored in red. In figures with IF, thicker outlines represent the cell perimeter and thinner outlines represent cell nucleus. A list of antibodies and fluorescent probes used for IF with their working concentration is shown below.

- ATF3. CST, #D2Y5W. 1/100.
- ATF4. CST. #D4B8. 1/100-1/200.
- JUN. CST, #60A8. 1/100-1/200.
- JUNB. Abcam, #245500. 1/100
- pJUN^Ser63^. CST, #2361. 1/100-1/200.
- FOSL1. Santa Cruz, #376148. 1/150.
- FOSL2. Santa Cruz, #166102. 1/150.
- IRF1. Abcam, #191032. 1/100.
- IRF7. Abcam, #115352. 1/100.
- PFKFB3. Proteintech, #13763-1-AP
- SLC2A1. Abcam #115730. 1/100.
- SOAT2. Santa Cruz, #59443. 1/50
- Phalloidin. Abcam, #176759. 1/1000.
- Donkey anti-mouse 488. ThermoFisher, #A21202. 1/200.
- Donkey anti-rabbit 555. ThermoFisher, #A31572. 1/200.

### Quantification of pixels intensity in ImageJ

ImageJ was used to quantify signal intensity of IF images. Briefly, three-colored images were split into single-colored images, nuclei was delineated using the freehand selection tool of ImageJ and intensity was calculated as the value of mean/area. Experiments were performed in duplicates. One representative experiment is shown.

### Immunofluorescence (IF) in frozen sections

Air pouch skin tissues were fixed in a 1:1 acetone:methanol mixture for 10 min at room temperature. Sections were blocked for 1 hr at room temperature in 3% BSA-PBS with 1% serum and 0.1% gelatin, and then incubated with primary antibody overnight at 4°C. After washing, Alexa Fluor secondary antibodies were added at 1:300 in 3% BSA-PBS for 1 hr, followed by 1:1000 DAPI for 20 minutes. After washing with PBS, slides were mounted with FluroSave. The antibodies and working concentrations used are listed below.

- F4/80. Invitrogen # MF48000 1/50
- SLC2A1. Santa Cruz #sc-33781 1/50
- p-JNK. Cell signaling #cs4668 1/50

### Immunohistochemistry in paraffin sections (IHC)

IHC analysis were performed using 3 μM sections of formalin-fixed paraffin-embedded samples of air pouch skin. After deparaffinization and rehydration, antigen retrieval was performed by boiling in citrate buffer pH 6 for 30 min. After antigen retrieval, endogenous peroxidase was inactivated with 3% H_2_0_2_ in water for 10 min at room temperature. Sections were incubated with 4% BSA-PBS for 1h at room temperature, and then with the specific primary antibody overnight at 4°C or 2h at room temperature. After washing, the Envision secondary reagent (DAKO) was added for 30 min at room temperature and sections were washed three times with PBS. 3,30-Diaminobenzidinetetrahydrochloride (DAB)(Vector, #SK-4100) was used as a chromogen. Sections were lightly counterstained with hematoxylin, dehydrated and then mounted. A non-related IgG was used as a negative control. The antibodies and working concentrations used are listed below.

- LDH. Santa Cruz #sc-33781 1/800
- ENO2. Proteintech #10149-1-AP 1/800
- iNOS. Abcam ab53004 1/2000

### Enzyme-linked immunosorbent assay (ELISA)

Protein levels in the supernatant of 250,000 cells stimulated with LPS or MSUc in triplicates, or in the lavage or serum from mice after MSUc injection in the air pouch were measured by ELISA (R&D Systems) following manufacturer’s specifications.

### Western blotting (WB)

For western blotting, proteins were extracted from 2 million cells unstimulated or stimulated with LPS or MSUc supplemented with protease inhibitors (Sigma Aldrich, #11697498001) Protein concentration was measured using the BCA reagent (Biorad, #500-0006). Proteins were resolved by standard SDS-PAGE gels and transfer to Immobilon PVDF membranes (Sigma, #IPVH00010) and incubated with specific antibodies overnight at 4°C. Membrane was then washed in PBS-Tween, incubated with Horseradish Peroxidase (HRP)-conjugated secondary antibodies and developed using Western Lighting Plus ECL reagent (Perkin, #0RT2655/0RT2755) via autoradiography (Genesee, 30-507). Densitometry analysis of digitalized western blotting images was performed using Fiji software (NIH). The antibodies and working concentrations used are listed below.

- SLC2A1. Santa Cruz #sc-7903 1/500
- PFKBP3. Proteintech #13763-1-AP 1/1000
- LDH. Santa Cruz #sc-33781 1/500
- TUBULIN. Cell Signaling, #3873. 1/1000.
- HPR-conjugated anti-mouse IgG. CST #7076. 1/2000.
- HPR-conjugated anti-rabbit IgG. CST #7074. 1/2000.

### Nuclear Magnetic Resonance (NMR) acquisition and processing

The metabolites from sera and supernatant were obtained by ultrafiltration (32) using a 3KDa filter (3kDa Omega, Pall Corporation, #OD003C34) with a final standard concentration of 0.5 millimolar (mM) in a total volume of 50 μL. The standard used was 3-(trimethylsilyl)propionic-2,2,3,3-d4 acid, sodium salt (TSP-d4) in deuterium oxide (Aldrich Chemistry, #293040-25G). From cell pellets, the polar metabolites were extracted by the precipitation method (33) with a final TSP-d4 concentration of 0.06617 mM. The one-dimension nuclear magnetic resonance (1D ^1^H-NMR) spectra were recorded using a 600 MHz Bruker Avance III NMR spectrometer fitted with a 1.7mm triple resonance cryoprobe. The standard Bruker pulse sequence “noesygppr1d” was used with a mixing time of 500 ms and 64 scans. A quality assurance procedure was performed before sample data acquisition, involving temperature checks and calibration as well as shim and water suppression quality. The data acquisition was obtained in the NMR facility of Skaggs School of Pharmacy and Pharmaceutical Science, University of California San Diego.

### Metabolites identification and quantification

The identification of metabolites approach was performed using the software Chenomx NMR suite 8.5 professional (Chenomx Inc., Edmonton, Canada) version 11, which contents a library of metabolites that match the peaks of compound according to their chemical shift. Metabolites concentration were normalized according to the standard TSP-d4 and the concentrations were reported in micromolar (μM).

### Mice

All animal experiments were performed in agreement with the Institutional Animal Care and Use Committee (IACUC). C57Bl/6 male mice (Jackson laboratories, Sacramento, CA, USA) were housed in a temperature-controlled room with a 06.00-18.00 hour light cycle and consumed regular chow and tap water ad libitum. Mice were 8-12 weeks at start of the experiments, and randomized before starting treatment to reduce bias.

### Air pouch model of inflammation

Dermal air pouches were established by injecting mice dorsally subcutaneously with 3 mL filtered (0.20 μm) air on day 0 and 3 (34, 35). On day 7, BAY-876 (5mg/kg, in 2.5%DMSO/ 40%PEG/ 5%Tween80) or JNK inhibitor (15mg/kg, in 2.5%DMSO/ 30%PEG/ 10%Tween80) were injected intrapouch and one hour later, MSUc were injected (3 mg in 0.5 ml PBS). 2.5%DMSO/ 40%PEG/ 5%Tween80 or 2.5%DMSO/ 30%PEG/ 10%Tween80 was used as vehicle control in mice injected with BAY-876 or JNKi, respectively. After eight hours of stimulation, pouch exudate was collected by rinsing with 1 mL endotoxin-free PBS, followed by 30 seconds of gentle massage. The collection was centrifuged (5 minutes, 450 × g) for analysis. Neutrophil quantification was perfomed after citospin and HE staining. Skin tissue covering the pouch was then excised and fixed in 10% formalin (Fisher #SF100-4) for subsequent haematoxylin and eosin (H&E) staining. Infiltrated cell area was scored using a qualitative score from 0 to 3 in a blinded manner for two independent readers. The number of mice used in each experiment is provided in the figure legends.

### Intra-articular MSUc injection

BAY-876 (7.5 mg/kg) or JNK inhibitor (15 mg/kg) were injected intraperitoneally one hour prior the injection of PBS in one knee or 100 μg of MSUc in the other knee (35). Eight hours after the injection of PBS or MSU, the knees were collected, fixed and decalcified (10% EDTA, Fisher #BP2482-500). Slides were stained with H&E. Infiltrated cell area was scored using a qualitative score from 0 to 3 in a blinded manner for two independent readers. Two cutting depths were used per knee and the largest infiltrated area per treatment was used. The number of joints per group are provided in the figure legends. 2.5%DMSO/ 40%PEG/ 5%Tween80 or 2.5%DMSO/ 30%PEG/ 10%Tween80 was used as vehicle control in mice injected with BAY-876 or JNKi, respectively

### Statistical analysis

Comparisons of quantitative data between groups was carried out using two-tailed Student T test unless otherwise indicated. Partial least-squares discriminant analysis (PLS-DA) was used to identify discriminant metabolites. Metabolites contributing to group discrimination in the PLS-DA were selected on the basis of having a variable important projection (VIP) score of >1. Box pots illustrate the median, Q1 and Q3 quartile of the data. Error bars in box plots represent the lowest and highest data point within 1.5x Q1-Q3 range. Bar graphs represent the media. Error bars in bar graphs represent the Standard error of the mean. All plots were generated using Numbers (iWORK’09), R studio (version 2.15.2 [2012-10-26]), and MetaboAnalyst version 5.0 (36). (#=P<0.10;*=P<0.05; **=P<0.01).

## SUPPLEMENTARY MATERIAL

**Supplementary Table 1**. Summary of RNA-Seq of human mDM treated with LPS or MSU for 5h.

**Supplementary Table 2**. Summary of RNA-Seq of mouse BMDM treated with LPS or MSU for 5h.

**Supplementary Table 3**. Summary of GSEA of RNA-Seq of human mDM treated with LPS or MSU for 5h.

**Supplementary Table 4**. Summary of GSEA of RNA-Seq of mouse BMDM treated with LPS or MSU for 5h.

**Supplementary Table 5**. List of primers used.

## AUTHOR CONTRIBUTION

Conceptualization: IC, AC, JMS, RLB, RT, ESL, CKG, MG

Formal Analysis: IC, AC, JMS, RC, ESL, RLB, MG

Investigation: IC, AC, JMS, RC, AT, AJL, JS, ESL

Data curation: IC, MG, JMS Original draft: IC, MG, CKG, RT

Writing-Review-Editing: IC, AC, MG, CKG, RT,

Funding acquisition: MG, RT, CKG

Visualization: IC, JMS, MG, CKG.

## Supplementary Figures

**Supplementary Figure 1.**
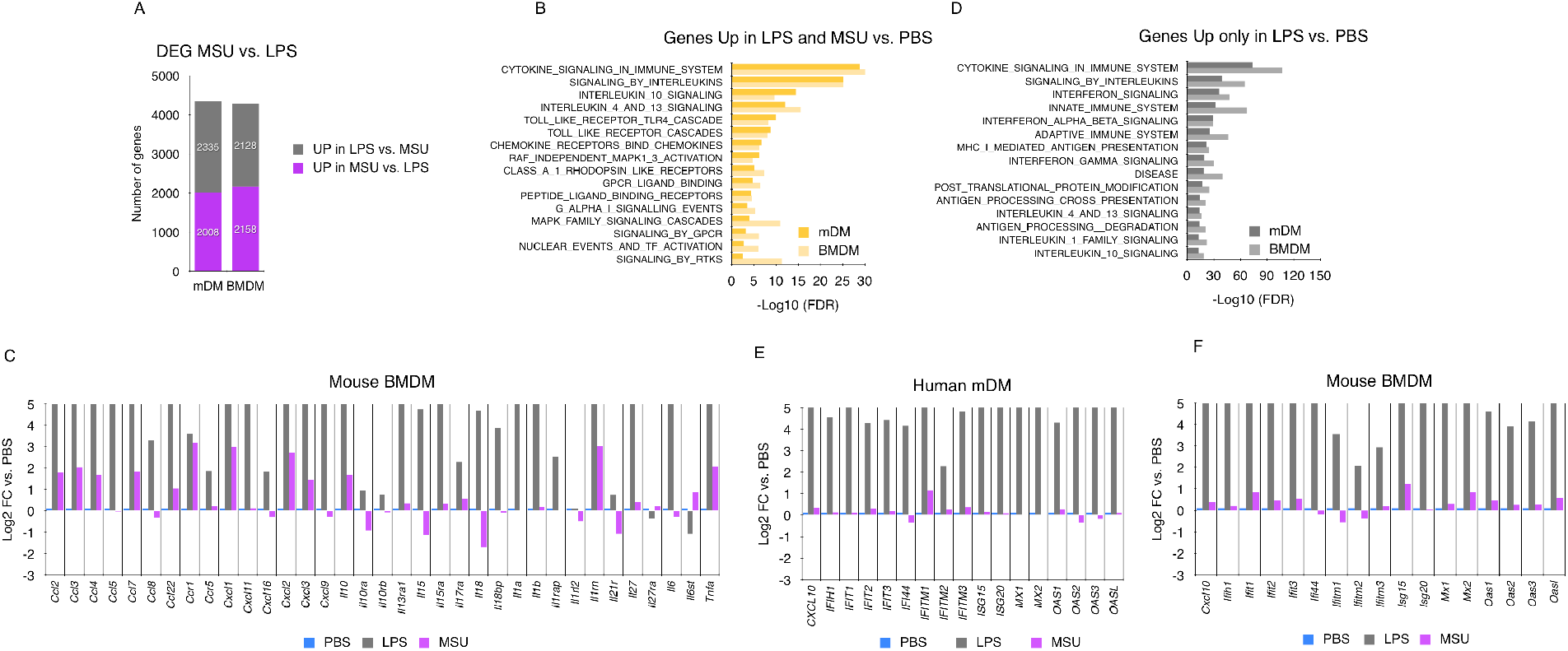
MSUc induces a unique transcriptional program with up-regulation of inflammatory cytokines but not interferon stimulated genes. (A) Bar plots showing similar number of genes up-regulated or down-regulated in mDM (left) or BMDM (right) treated with MSUc vs. LPS. (B,C) GSEA analysis using REACTOME of genes up-regulated in LPS vs. MSUc (B) or uniquely up-regulated in LPS (C) in BMDM and mDM showing enrichment in inflammatory pathways including response to interferon. (D) Bar graphs showing the up-regulation of several inflammatory cytokines by LPS and MSUc with higher degree of up-regulation in LPS. (E,F) Bar graphs showing induction of the interferon program in BMDM (E) or mDM (F) treated with LPS but not MSUc.

**Supplementary Figure 2.**
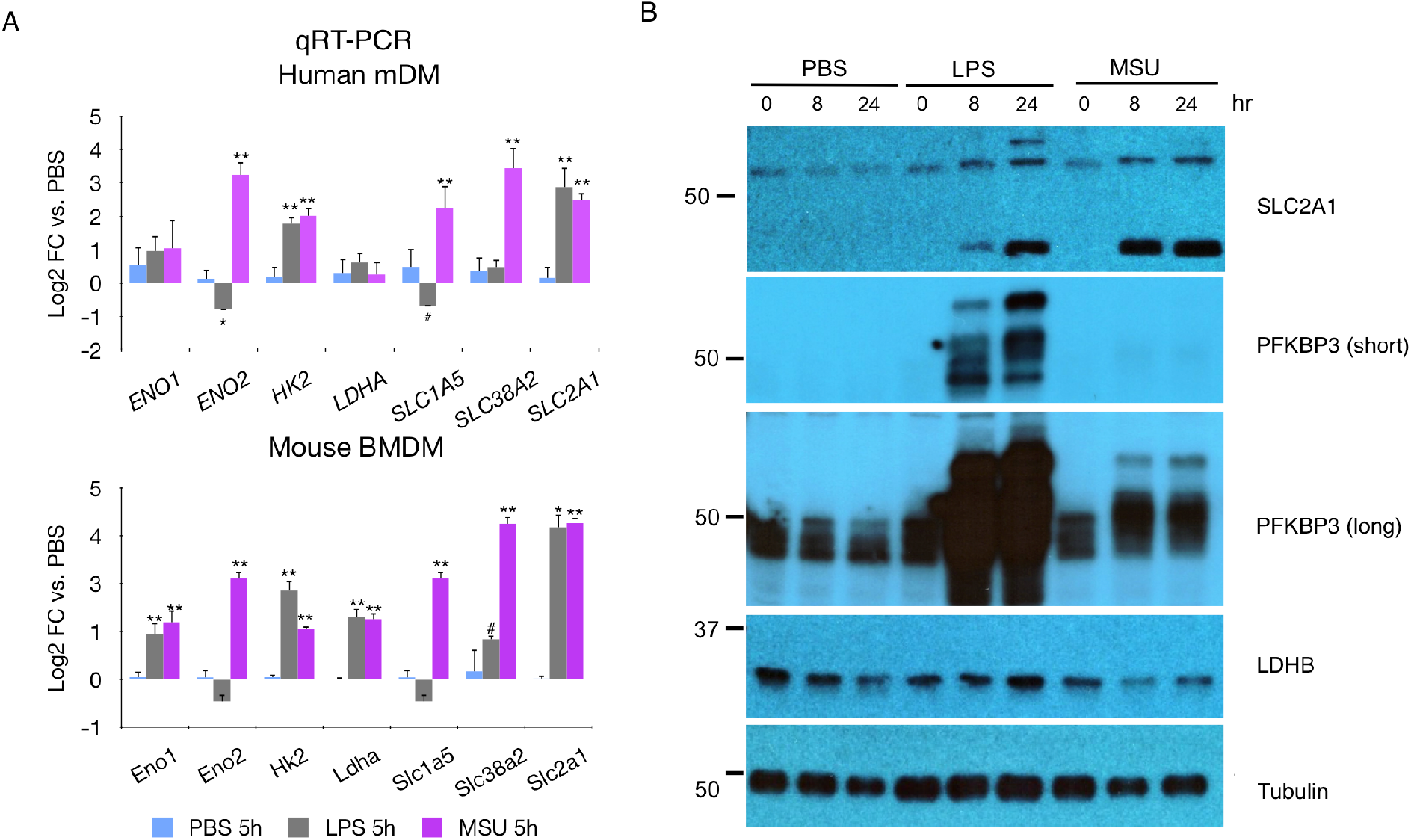
Treatment with MSUc leads to an altered metabolic program. (A) RT-qPCR showing up-regulation of metabolic genes in mDM (above) or BMDM (below) treated with LPS or MSUc for 5h (n≥4/condition). (B) WB of lysates of BMDM treated with LPS or MSUc in control PBS, LPS or MSUc at 8h or 24h showing up-regulation of SLC2A1 and PFKBP3 in BMDM treated with LPS or MSUc and down-regulation of LDHB in BMDM treated with MSUc.

**Supplementary Figure 3.**
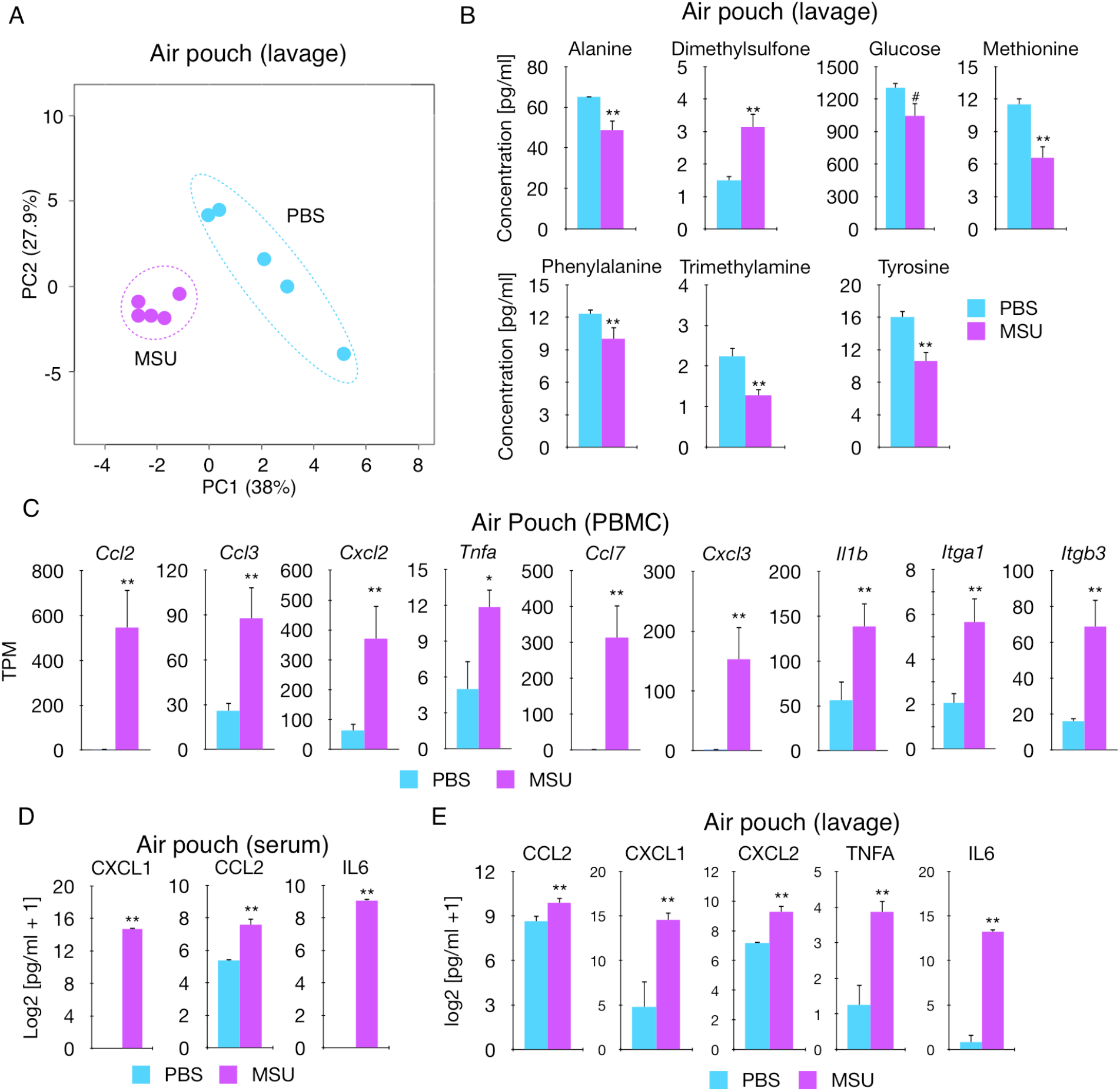
Injection of MSUc in the air pouch leads to systemic alteration of inflammatory and metabolic profile. (A) PLS-DA graph of levels of metabolites in the air pouch lavage of mice injected with PBS or MSUc as assessed by 1D ^1^H-NMR showing the divergence between groups (n=5 mice/group). (B) Bar graphs showing reduced concentration of alanine, glucose, methionine, phenylalanine, trimethylamine, and tyrosine, and increase of dimethylsulfone in the air pouch lavage of mice injected with MSUc. (C) Bar graphs showing expression analysis by RNA-Seq (TPM) of inflammatory genes up-regulated in PBMC from mice injected with MSUc in the air pouch. (P<0.05). (D,E) CXCL1, CCL2 and IL6 levels in the serum (D) and additionally CXCL2 and TNFA levels in the air pouch lavage (E) of mice injected with MSUc in the air pouch assessed by ELISA (n=5 mice/group). (#=P<0.10;*=P<0.05; **=P<0.01).

**Supplementary Figure 4.**
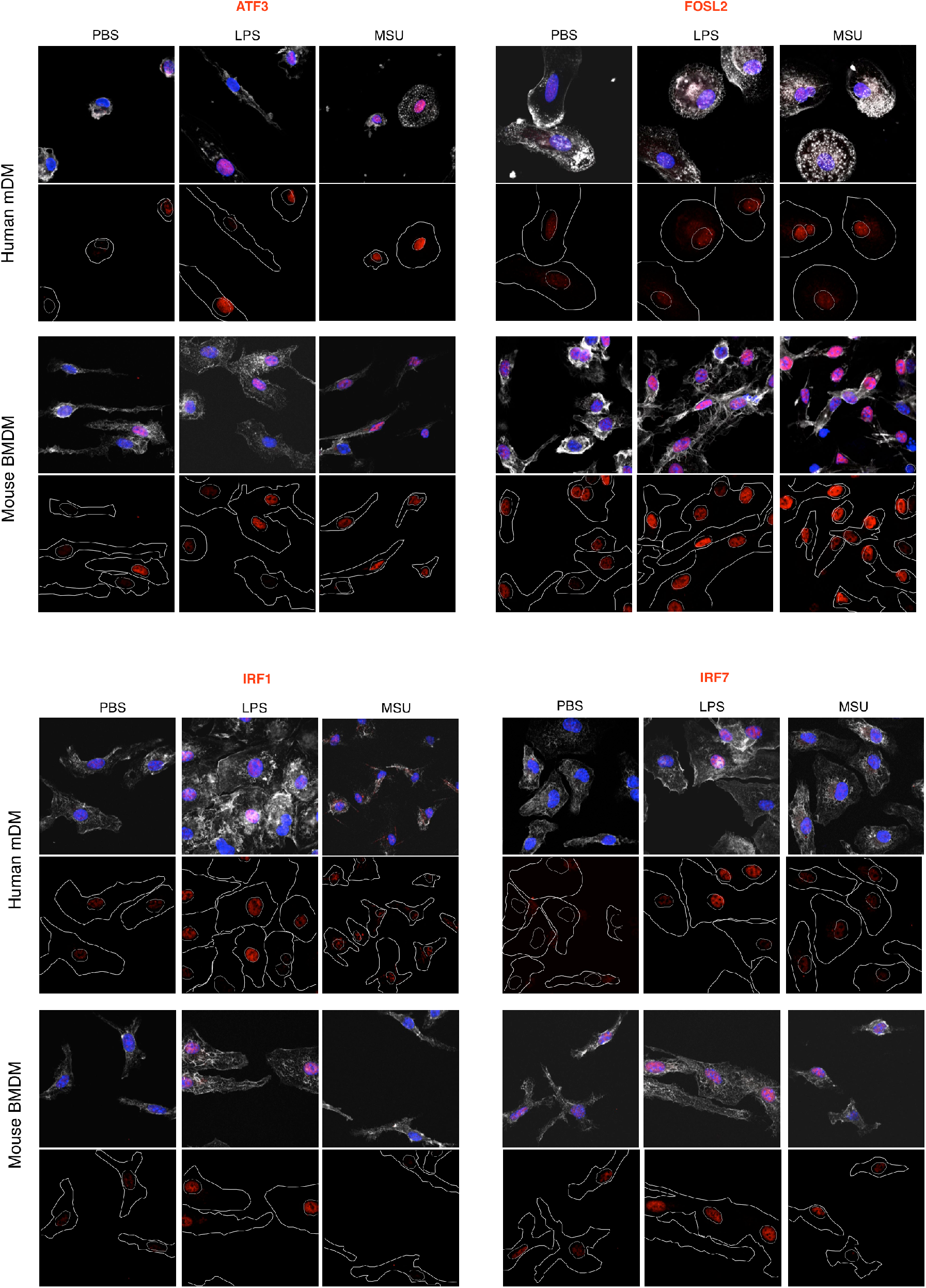
MSUc leads to up-regulation of AP-1 proteins but not IRF1 or IRF7. Protein analysis by IF of mDM and BMDM treated with LPS or MSUc for 5h showing up-regulation of AP-1 proteins by MSUc and LPS but up-regulation of IRF1 and IRF7 only by LPS. (n=3/condition).

**Supplementary Figure 5.**
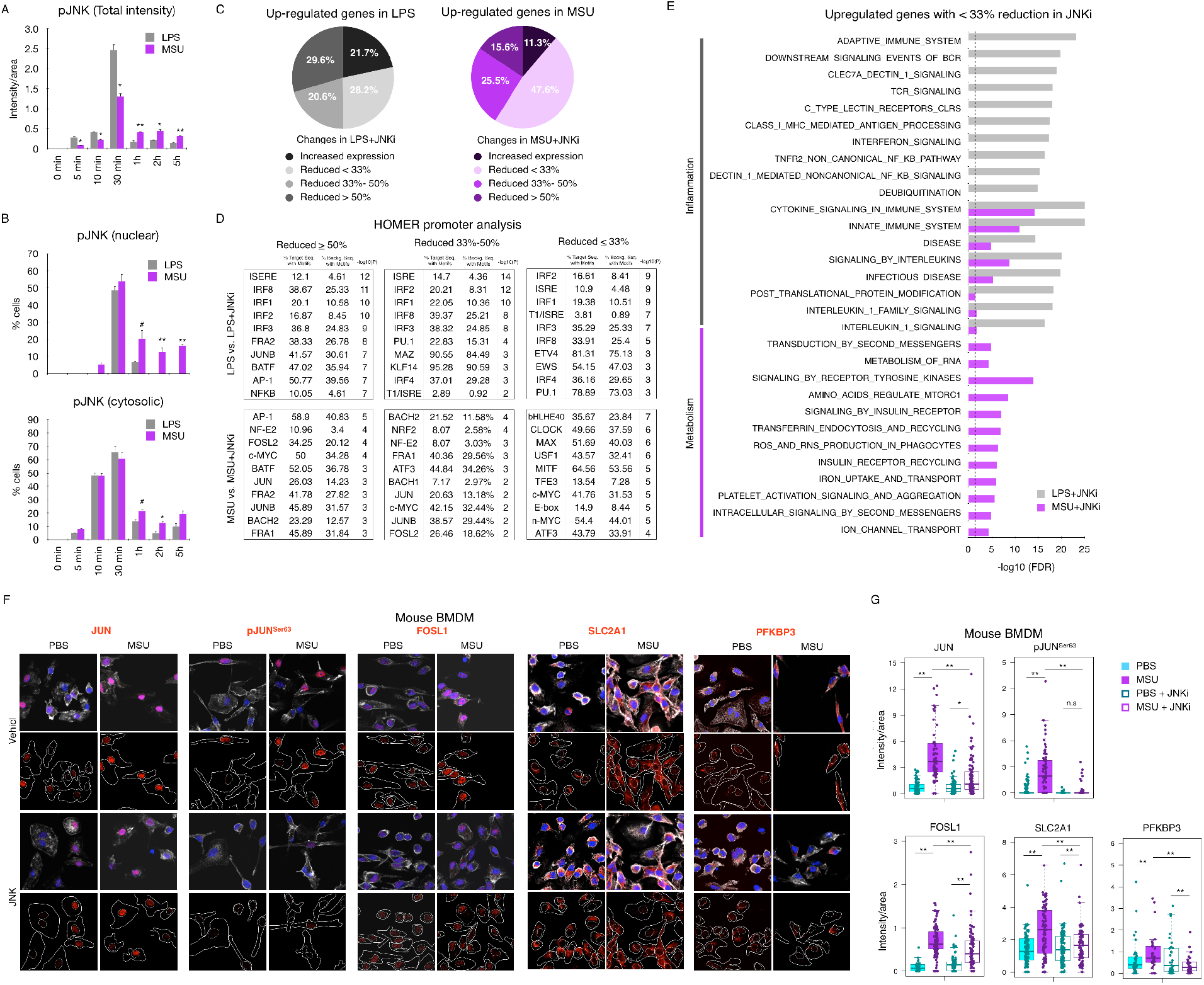
JNK activation by MSUc is required for up-regulation of inflammatory and metabolic genes. (A,B) Quantification of pJNK expression in BMDM treated with LPS or MSUc at various time points showing increased total pJNK expression in BMDM treated with LPS for 30 min (A) but prolonged pJNK expression in BMDM treated with MSUc (A,B)(n=2 replicates/condition). (C) Pie chart showing the degree of amelioration of gene expression in BMDM treated with LPS+JNKi vs. LPS or MSUc+JNKi vs. MSUc for 5h. (D) HOMER analysis of the promoter [-2000;+500 bp, TSS] of genes up-regulated by LPS or MSUc and further down-regulated by treatment with JNKi (>50% left, 33%-50% center, <33% right). Data shows enrichment in IRFs motifs in genes ameliorated by JNKi in LPS and motifs for AP-1, MYC, NRF2 and circadian clock proteins in genes ameliorated by JNKi in MSUc. (E) GSEA analysis using REACTOME of genes between > 50% reduccion of expression in MSUc+JNKi vs. MSUc or LPS+JNKi vs. LPS. Data shows enrichment in inflammatory gene sets in LPS and MSUc and metabolic gene sets in MSUc. (F,G) Protein analysis by IF of JUN, pJUN^Ser63^, FOSL1, SLC2A1 and PFKBP3 in BMDM treated with MSUc or MSUc+JNKi for 5h. Quantification of G corresponds to experiment shown in F and shows downregulation of protein expression in MSUc+JNKi vs. MSUc (n=3/condition). (#=P<0.10;*=P<0.05; **=P<0.01). Broken lines in E represents the cutoff for significance –log_10_(0.05)=1.30.

**Supplementary Figure 6.**
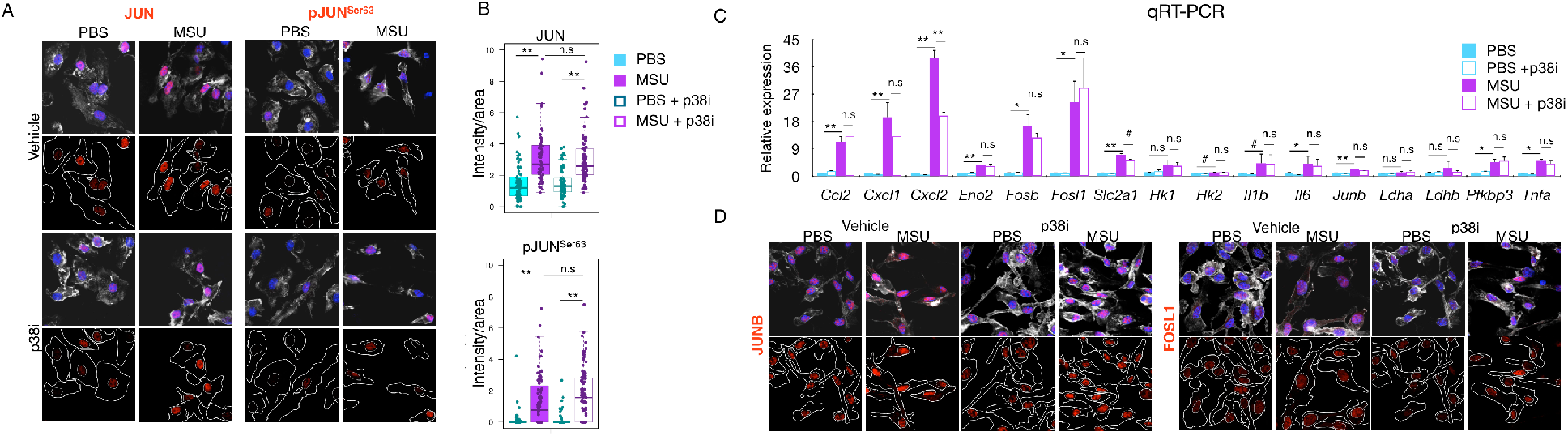
p38 is not required neither for the up-regulation of JUN and pJUN^Ser63^ nor the overexpression of inflammatory and metabolic genes. (A,B) Protein analysis by IF showing undetectable changes in levels of JUN and pJUN^Ser63^ in BMDM treated with MSU+p38i vs. MSUc for 5h. (C) RT-qPCR analysis of inflammatory and metabolic genes showing undetectable changes –except for *Cxcl2*-after treatment with p38i. (D) Protein analysis by IF showing undetectable changes of JUNB and FOSL1 after treatment with p38i (n=2/condition). (#=P<0.10;*=P<0.05; **=P<0.01).

**Supplementary Figure 7.**
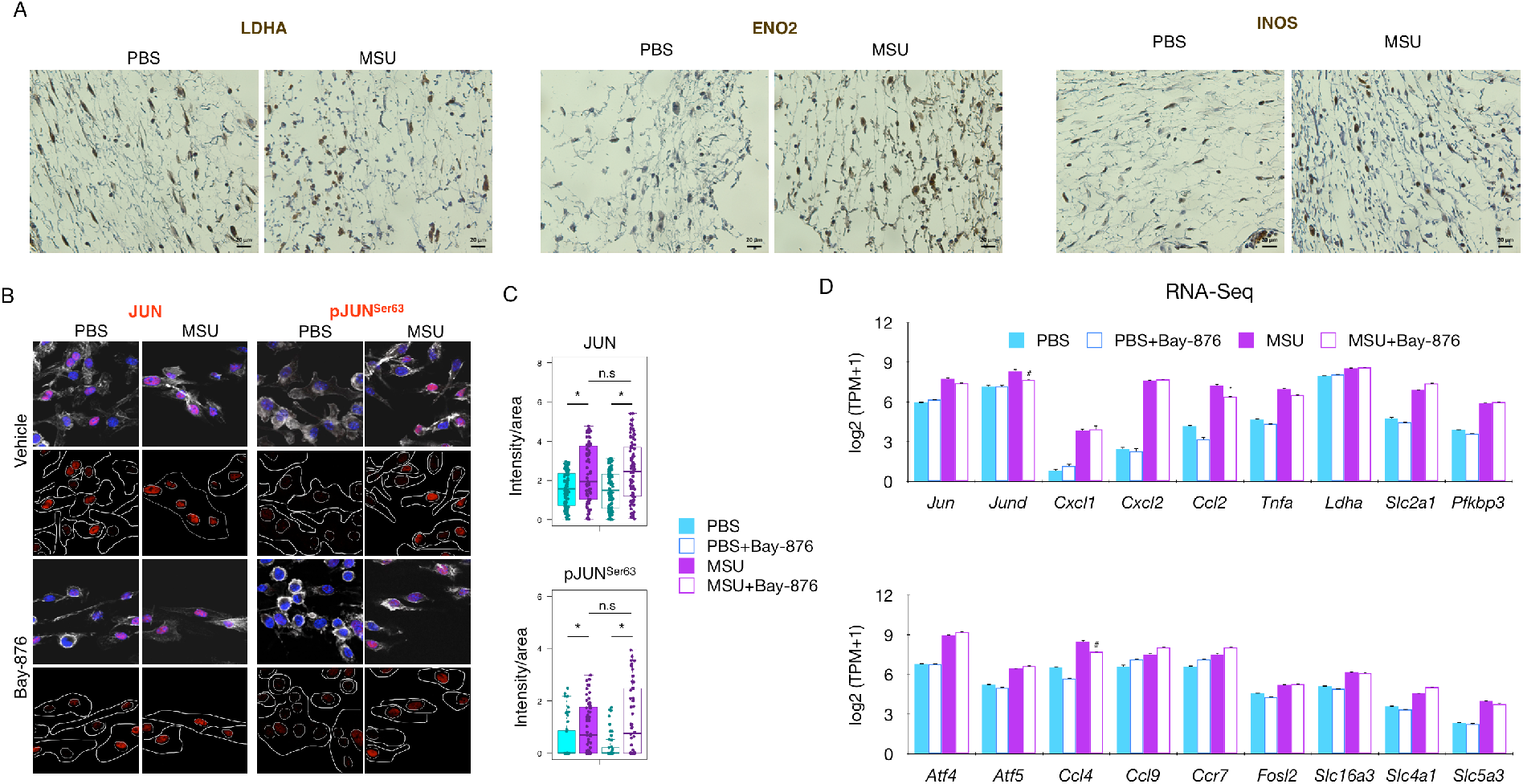
MSUc leads to up-regulation of glucose metabolism related enzymes in macrophages of the air pouch. Blockage of SLC2A1 signaling does not alter the up-regulation of JUN, pJUN^Ser63^, inflammatory or metabolic genes by MSUc *in vitro*. (A) Protein analysis by IHC of LDHA, ENO2 and iNOS in histological sections of air pouch injected with PBS or MSUc (n=3/condition). (B,C) Protein analysis by IF in BMDM treated with MSUc or MSUc+Bay-876 for 5h showing undetectable changes of JUN and pJUN^Ser63^ (n=3/condition). (D) Expression analysis by RNA-Seq (TPM) of BMDM treated with MSUc or MSUc+Bay-876 for 5h showing undetectable or minor changes in the expression of inflammatory and metabolic genes after treatment with Bay-876 (n=5/condition). (#=P<0.10;*=P<0.05; **=P<0.01).

